# Microbial community dynamics during a harmful *Chrysochromulina leadbeateri* bloom in northern Norway

**DOI:** 10.1101/2022.06.21.496960

**Authors:** Nerea J. Aalto, Hannah Schweitzer, Erlend Grann-Meyer, Stina Krsmanovic, Jon B. Svenning, Lars Dalheim, Sebastian Petters, Richard Ingebrigtsen, Chris J. Hulatt, Hans C. Bernstein

## Abstract

A harmful algae bloom occurred in late spring 2019 across multiple, interconnected fjords and bays in northern Norway. The event was caused by the haptophyte *Chrysochromulina leadbeateri* and led to severe fish mortality at several salmon aquaculture facilities. This study reports on the spatial and temporal succession dynamics of the holistic marine microbiome associated with this bloom by relating all detectable 18S and 16S rRNA gene ASVs to the relative abundance of the *C. leadbeateri* focal taxon. A k-medoids clustering enabled inferences on how the causative focal taxon co-bloomed with diverse groups of bacteria and microeukaryotes. These co-blooming patterns showed high temporal variability and were distinct between two geographically separated time series stations during the regional harmful algae bloom. The distinct blooming patterns observed with respect to each station were poorly connected to environmental conditions suggesting that other factors, such as biological interactions, may be at least as important in shaping the dynamics of this type of harmful algae bloom. A deeper understanding of microbiome succession patterns during these rare but destructive events will help guide future efforts to forecast deviations from the natural bloom cycles of the northern Norwegian coastal marine ecosystems that are home to intensive aquaculture activities.

## INTRODUCTION

Harmful algal blooms (HABs) are common in warm and temperate waters, but also occur in northern cold-water ecosystems (Hallegraeff et al., 2021; Karlson et al., 2021). These blooms pose a serious risk for aquaculture activities, because farmed fish cannot escape from developing adverse conditions caused by marine microalgae. Previously, the most infamous northern Norwegian bloom occurred in 1991 in Ofotfjord causing high mortality in affected fish farms (Rey, 1991). Smaller HAB incidents have occurred since then, although they have gained little mainstream attention (Grann-Meyer, 2020). The salmon aquaculture industry has expanded tremendously over the past decades to a landing value of salmon that reached 7 billion USD in 2019 (Statistics Norway, 2020). Therefore, the Norwegian economy and seafood export supply was seriously threatened when a deadly HAB of *Chrysochromulina leadbeateri* (haptophyte) reoccurred and caused massive fish mortalities in the Troms and Nordland regions of costal northern Norway from mid-May to the beginning of June 2019 (Karlsen et al., 2019). The fish loss was ca. 14 500 tonnes, which is the highest amount among fish kill related HABs in Scandinavia since 1988 (Karlsen et al., 2019; Karlson et al., 2021). Knowledge that can eventually inform causation is very limited, especially in the context of seasonal microbial blooming dynamics, detailed taxonomy of co-blooming species, and the environmental conditions during the 2019 incident.

Marine HABs are often associated with altered ratios of inorganic nitrogen-to-phosphate (N:P), coupled with a long period of high irradiance, calm weather, and strong surface water stratification (Nielsen et al., 1990; Anderson et al., 2002; Uronen et al., 2005). However, the physical environment is not the only determining factor for the strength and duration of a HAB (Davidson et al., 2012) as more evidence is showing that biological interactions impact microalgal bloom dynamics. The roles and importance of co-blooming members of the marine microbiota is becoming a more recognized factor in structuring the temporal and spatial relationships of phytoplankton communities (e.g., Needham and Fuhrman, 2016; Martin-Platero et al., 2018; Aalto et al., 2022). Therefore, it is important (yet uncommon today) to observe and catalogue the blooming patterns of whole surface water microbiome while studying the ecology of focal taxa involved with HAB events.

Time series analysis combined with high-resolution taxonomic screening such as 16S and 18S ribosomal RNA gene amplicon sequencing variants is an important method for evaluating microbial succession dynamics and the ecological factors that drive blooms (Fuhrman et al., 2015; Martin-Platero et al., 2018). However, knowledge about the nature of taxon-specific and non-specific blooming patterns is poor because identification of these “hidden” dynamics is challenging. Therefore, an increasing number of modern analytical tools has been developed to apply with time series analysis. For example, WaveClust (Martin-Platero et al., 2018) and k-medoid (Kaufman and Rousseeuw, 2009; Coenen et al., 2020) are applicable and promising approaches to infer both temporal dynamics and networks of microbial interactions.

The spring-summer phytoplankton seasonal succession along the coast of northern Norway has been extensively studied but the majority of data collected to-date has used only morphology-based identification of microalgal taxa, which limited the resolution of taxonomic diversity of these blooming patterns (e.g., Eilertsen et al., 1981; Reigstad and Wassmann, 1996; Degerlund and Eilertsen, 2010; Aalto et al., 2021). From previous studies, we know that the onset of the “spring bloom” occurs in late March/early April, and it is driven by increased irradiance as the region undergoes a shift from polar night to spring, within elevated winter nutrient concentrations and a well-mixed water column (Eilertsen and Taasen, 1984; Eilertsen and Frantzen, 2007; Aalto et al., 2021). These conditions lead to a rapid and intense increase of many chain forming centric diatoms that co-bloom with *Phaeocystis pouchetii* (haptophyte), which often outlasts the diatoms and persists throughout the summer (Eilertsen and Taasen, 1984; Degerlund and Eilertsen, 2010). The spatial and interannual variation of the late bloom (May-June) phytoplankton community can comprise a diverse mix of low-nutrient-tolerant, heterotrophic, and mixotrophic species belonging to dinoflagellates, cryptophytes, ciliates, and other small flagellates (Eilertsen et al., 1981; Aalto et al., 2021). The roles and taxonomic succession patterns of bacterioplankton in these annual blooms is essentially unknown for the region.

Our aim was to contribute to the knowledge of the ecology and microbial succession dynamics associated with the *C. leadbeateri* HAB which role as a causative agent for this devastating event has already been established (Karlsen et al., 2019; John et al., 2022). This was accomplished by asking three targeted research questions: i) how did the marine microbiome composition change during the 2019 HAB event in relative relation to the focal taxon (*C. leadbeateri*)?; (ii) what were the major blooming (temporal) patterns among prokaryotes and microeukaryotes and which taxa co-bloomed with the focal taxon?; and iii) which environmental factors were associated with the observable differences in the marine microbiome composition? This study provides a molecular-based investigation of the destructive, 2019 *C. leadbeateri* HAB and presents new insights into a seasonal marine microbial ecosystem that can have a major impact on the social-economic welfare of Norway through its relation to the aquaculture industry.

## MATERIALS AND METHODS

### Study areas and sample collection

Samples for microbial community analyses were collected from two primary time series stations and in addition from eight single time point stations. Time series sampling was performed between May 27 (sample day 0) and June 21 (sample day 25) in 2019 (Supplementary Table S1). The first station, at the vicinity of a fish farm (Lerøy Aurora at Solheim) was named Grøtsund sampling station (**GSSS**, 69.80 N, 19.36 E). The second station, named as Finnfjord sampling station (**FSS**, 69.19 N, 18.03 E), is located 60 km south of GSSS, within the same interconnected fjord-sound system (Figure 1a). These time series stations were chosen because they present historical sample locations (Aalto et al., 2021) and provided long-term access to sampling facilities (vessel: ∼5 m Polarcirkel and laboratories). The single time point stations were chosen to gain a geographical representation of the bloom, given that it was infeasible to run time series collections across the entire affected region.

**Figure 1.**
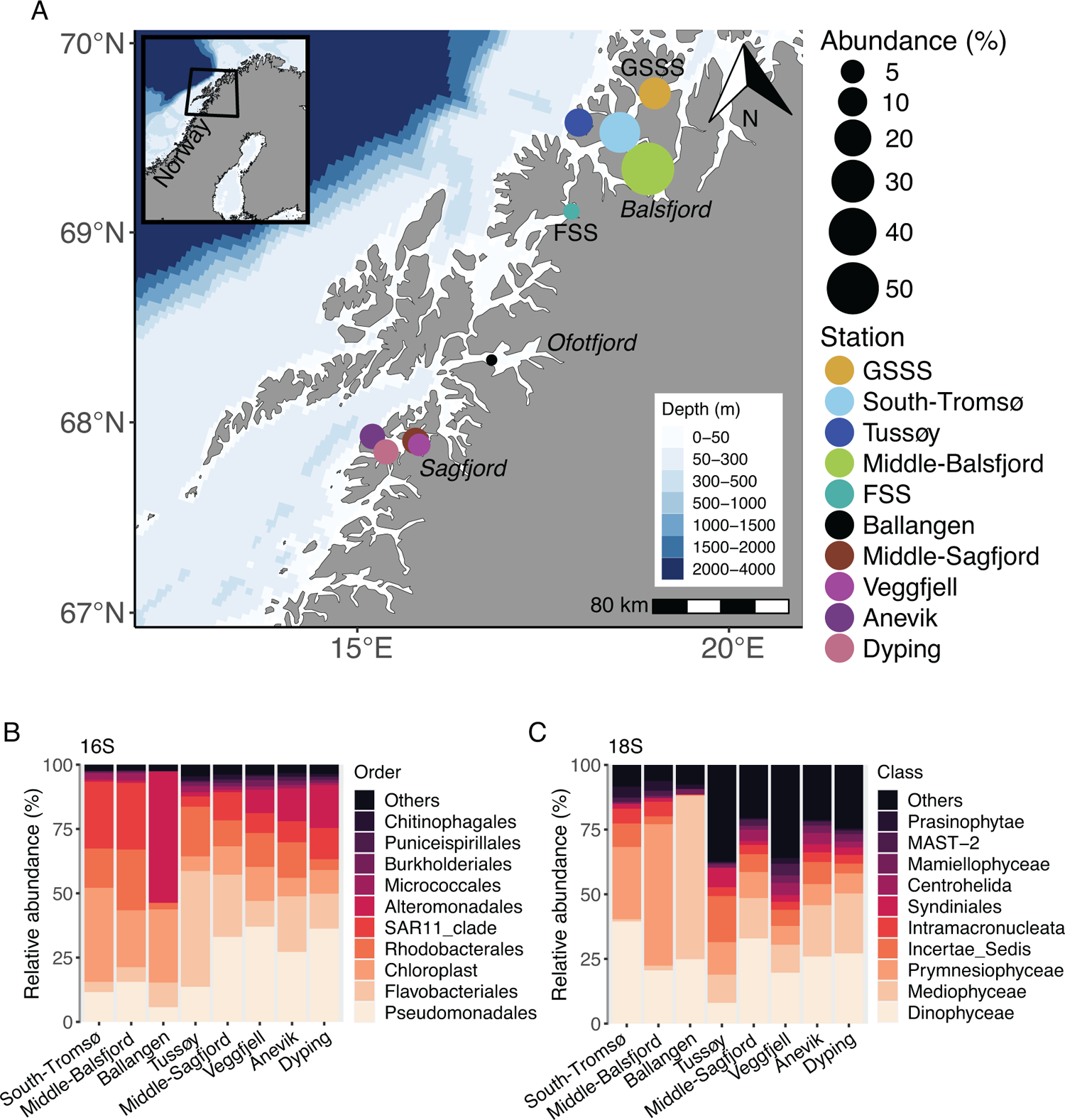
A map showing the geographical area in northern Norway where *Chrysochromulina leadbeateri*-related harmful algal bloom occurred in summer 2019. The main sample locations are *named* on the map. **A)** Microbial DNA for 18S rRNA and 16S rRNA amplicon sequence analysis was collected once from the marked stations, except from FSS (Finnfjord sampling station) and GSSS (Grøtsund sampling station) which are the primary stations of this time series study and were sampled 7 and 8 times, respectively, between May 27^th^ and June 21^st^. The size of the circles represents the relative abundance of the *C. leadbeateri* focal taxon (average relative abundance for FSS and GSSS stations across all samples). Not all of the stations represent localities where HAB-related fish mortality was experienced. Taxonomic composition of the most abundant **B)** prokaryotes (16S) and **C)** microeukaryotes (18S) retrieved from Amplicon Sequence Variants (ASVs) tables specified at the order and class taxonomic rank level, respectively. Note that FSS and GSSS stations are excluded from **B** and **C** as the taxonomic composition across time series is shown in Figure 2.

Samples were collected from each time series station twice a week, except station FSS was sampled once in the second sampling week (starting the week of May 27, 2019), from fixed sampling depth of 10 m using a 1.7 L Niskin bottle to get approximately 8.5 L of seawater. The collected seawater was prefiltered for large biomass with a 150 μm plankton mesh and transported to shore in a 15 L Nalgene plastic carboy. All sampling equipment was rinsed with 2% bleach prior and between each sampling effort followed by a rinsing with MilliQ water and a minimum of two volumes of station sampling water prior to collection. Temperature and salinity profiles down to 15–50 m were recorded with a hand-held AML oceanographic X2 electron conductivity-temperature-depth (CTD) instrument (AML Oceanographic, Canada). Seawater particles – i.e., microbial biomass – was collected by vacuum filtration onto 0.22 μm polycarbonate filters (GE Healthcare Whatman 47 mm Nucleopore Polycarbonate Track Etched Membranes; GE Healthcare 111106). All filtering was done inside a sterile laminar flow hood, and the total filtered volume per biological replicate (n = 6) varied between 500-750 mL. Filters were immediately flash frozen and stored at −80 °C. All filtering equipment was disinfected with 2% bleach prior and between sampling efforts and rinsed with MilliQ water. Water samples for macronutrients: silicate (Si(OH)_4_), phosphate (PO_4_^3-^), and nitrate + nitrite (NO_3_^-^ + NO_2_^-^), were filtered through 47 mm GF/F glass microfiber filters (Whatman) into Falcon tubes using disposable syringes and stored at −20 °C.

Filtered microbial biomass was collected from the eight single time point stations between May 27^th^ and June 21^st^ (Figure 1a and Supplementary Table S1). No additional environmental data was collected from these stations. The data of these samples (n = 3 or n = 6 per station) is not included within time series analyses but used to compare the 2019 HAB affected area via spatial variation in the relative abundance of the focal taxon and associated microbial communities. The DNA extraction and downstream analysis of all samples were performed at the UiT – The Arctic University of Norway as described below.

### Environmental factors

The vertical surface water properties from the time series sampling stations (FSS and GSSS) were inspected using CTD-profile data (Supplementary Figure S1 and S2). The difference in surface water stratification between sampling stations was determined by calculating the mixed layer depth (MLD) from each CTD-profile data using potential density. The depth of 1 m was used as a reference depth as the deployed handheld CTD-sensor enables measurements near surface. A density threshold of 0.1 kg m^-3^ was used to define MLD – i.e. the depth below 1 m where density first exceed the defined threshold (Peralta-Ferriz and Woodgate, 2015). Temperature and salinity values from the depth of 10 m (corresponding the depth of collected biomass) as well as MLD values were further used in statistical analyses. As the CTD-casts between sampling days 11 and 17 were unsuccessful, temperature, salinity, and MLD values for these days were interpolated using the R function ‘na.approx’ from the R package ‘zoo’ to fill the missing values. Nutrient concentrations were analyzed 20 months later at the UiT – The Arctic University of Norway with a Quaatro39 autoanalyser (Seal Analytical, UK). PO_4_^3-^ concentrations below the detection limit of 0.04 µmol L^-1^ were replaced with a small value (0.001) to enable the estimation of this variable in statistical analysis.

### Unialgal control cultures

We verified the focal taxon within field samples using unialgal control that was performed with the *Chrysochromulina leadbeateri* UiO-035 (Edvardsen et al., 2011) culture obtained from the Norwegian Culture Collection of Algae (NORCCA) repository and cultivated at 6 °C on f/2 medium (Guillard, 1975), fortified with 20 nM of selenium. The cultured *C. leadbeateri* was isolated from the toxic bloom of 1991 in Vestfjorden, northern Norway (Eikrem and Throndsen, 1998). Taxonomic identification of the cultured isolate has been mainly based on scale morphology in comparison to holotype (Estep, 1984) and other previous reports on *C. leadbeateri*-like cells both from northern and southern hemisphere (Eikrem and Throndsen, 1998). Several *Chrysochromulina* species are characterized by species-specific bilayer scales which have been considered a morphotype-specific character (Eikrem and Throndsen, 1998). The near-full length 18S SSU rRNA gene of the *C. leadbeateri* UiO-035 has been sequenced and is found in Genbank under the accession number AM491017. Cells were collected (n = 6) and processed identically to the field samples described below.

### Amplicon sequencing

Genomic DNA was extracted and downstream amplicon sequences were obtained using the same protocol as described in Aalto et al. (2022). All the samples were processed together in accordance with the Earth Microbiome Project protocols (Gilbert et al., 2010) with modifications according to Amaral-Zettler et al. (2009). This included recommended primers and barcodes for the V4 hypervariable region of the 16S SSU rRNA gene using the V4 forward (515F) and V4 reverse (806R) primers, and the V9 hypervariable region of the 18S SSU rRNA was targeted with the V9 forward (1391F) and V9 reverse (EukBr) primers. Biological replicates were sequenced on an Illumina MiSeq instrument (Illumina, San Diego, CA) according to (Caporaso et al., 2010) at Argonne National Laboratory (Lemont, IL, USA). The realized length of forward and reverse reads for 18S data set varied between 151 and 282 bp and for 16S data set the corresponding range was 152-289 bp.

### Amplicon analysis

The amplicon analyses were performed through the QIIME2 environment and using QIIME2 plugins as previously described (Aalto et al. 2022). Briefly, Illumina reads and the corresponding barcode files were imported and demultiplexed using the Earth Microbiome Project paired end flag. All reads were filtered, de-replicated, and chimera checked using all default parameters in the DADA2 v2021.2.0 (Callahan et al., 2016). Reads were merged and amplicon sequence variants (ASVs) were determined using DADA2 v2021.2.0. The DADA2 statistic on sequence reads is provided in Supplementary Data S1. A 16S and 18S rRNA gene classifier from the SILVA v138.1 database was trained using RESCRIPt (Quast et al., 2012; Robeson et al., 2020). The ASVs were classified with the self-trained classifier database (Yilmaz et al., 2014). It is noted that the major challenge in 16S and 18S based taxonomy is its dependency on taxonomic resolution limitations of the database. We chose to use all classifications given from the SILVA v138.1 database because it is a standardized approach. Manual curation of taxonomic classification except unialgal controls was beyond the scope of this study.

Within the unialgal control cultures the 18S amplicon analysis assigned four abundant ASVs (Supplementary Figure S3a). All four of these ASVs had identical taxonomic assignments within the SILVA v138.1 database: Family = Prymnesiales, Genus = OLI16029, Species = Unknown. Of these four ASVs assigned as OLI16029, ASV *24a92740c5af4cd5a5ac0db70830fbc0* (hereafter referred to as the “focal ASV”) was the most abundant and comprised 88% of the sequence reads (Supplementary Figure S3b). All four of these ASVs were also present among field samples with seven additional ASVs assigned as OLI16029. The sequence of the focal ASV was manually searched against NCBI BLAST. The results showed sequence match of 100 % (Query Cover 91 %) with *C. leadbeateri* UiO-035 but also 99.40 % identity (Query Cover 97 %) was found with *C. leadbeateri* UiO-393 strain (NORCCA; Genbank accession number ON815372), which was isolated during 2019 HAB by John et al. (2022). The sequence alignment of all ASVs classified indicative to OLI16029 as well as *C. leadbeateri* strains of UiO-035 and UiO-393 are shown in Supplementary Figure S4. Hereafter these ASVs are referred as *C. leadbeateri*.

### Statistics, clustering and correlation

All ASVs not assigned to the expected kingdom (*Archaea*, *Bacteria* and *Chloroplast*) were removed along with all mitochondria assignments from 16S data set. ASVs assigned to *Archaea*, *Bacteria*, *Vertebrata* and *Arthropoda* and phyla *Cnidaria*, *Echninodermata*, *Annelida*, *Nematozoa*, *Nemertea*, *Porifera*, *Mollusca*, *Ctenophore* and *Tunicata* or phyla belonging to macroalgae and land plants or land fungi such as *Bangiales*, *Floridephycidae*, *Ochrophyta*, *Phragmoplastophyta*, *Ascomycota*, *Basidiomycota* and *Myxogastria* were removed from the 18S data set. Additional information on number of ASVs and sequencing depths before and after removing taxa is provided in Supplementary Table S2. The total number of classified ASVs in FSS for 18S and 16S data set was 900 and 1309, respectively, and in GSSS 2196 and 1865, respectively. As an exception regarding correction of taxonomic annotations, classified 16S order *Enterobacterales* was manually changed to closely related order *Alteromondales* as the main genera of this order were *Pseudoalteromonas*, *Colwellia* and *Glaciecola* that are known to be members of *Alteromonadales* (Parte et al., 2020).

Downstream analysis was completed in R (R Core Team, 2021), using the ‘microeco’ (Liu et al., 2021) and ‘vegan’ packages (Oksanen et al., 2013). This study also made use of the K-medoids clustering method adapted from a previously described approach applied to time series microbiome data to examine distinct temporal trends in blooming patterns within communities (Coenen et al., 2020). Briefly, ASV counts from each sample type – i.e., sampling station (FSS or GSSS) and PCR amplification primer type (16S or 18S) – were preprocessed via z-score and variance stabilizing transformations using the ‘DESeq2’ package in R (Love et al., 2014). Pairwise distance matrices were calculated by the Euclidean distance prior to partitioning each sample specific ASV into k-medoid clusters according to similarity criterion over the time series. The quality of each cluster per each sample type was assessed via the Calinski-Harabasz index (Supplementary Figure S5) (Lord et al., 2017). Pearson’s correlation was used to examine the relationship between each temporal dynamic, represented by medoid ASV (cluster centroids), and environmental measurements but also to obtain the positively and negatively co-blooming microalgae and bacterial taxa with focal ASV.

The difference in microbial communities, i.e. beta-diversity, between sampling stations was measured using unweighted UniFrac distance metric which accounts occurrence – presence/absence of ASV – and phylogenetic diversity (Lozupone et al., 2011). The beta-diversity and relationship between community compositions and environmental factors (temperature, salinity, MLD, Si(OH)_4_, PO_4_^3-^, NO_3_^-^ + NO_2_^-^, and relative abundance of focal taxon (*C. leadbeateri*)) was visualized and conducted via distance based redundancy analysis (dbRDA). The permutational multivariate analysis of variance (PERMANOVA) was performed to test the (dis)similarity in prokaryotic and microeukaryotic community compositions between stations (pairwise) and within stations (Anderson, 2001). The Tukey test was performed on inorganic nutrients to determine if the mean concentration values were statistically different between stations (Abdi and Williams, 2010).

### Data repository and reproducible analyses

All sequencing including raw fastq files and environmental data is available on the Open Science Framework (osf.io), along with all R Markdown scripts used for analyses and graphing (osf.io/4wjhp/). All Supplementary data S1-S11, referred to in this study, is available and found on osf.io/4wjhp/.

## RESULTS

### Geographical distribution of focal taxon in relation to fish mortality

We detected the *Chrysochromulina leadbeateri* focal taxon in varying relative abundance across the bloom-affected region of the northern Norwegian coast and fjords (Figure 1a). This data is in concordance with other investigations of this 2019 HAB event that established *C. leadbeateri* as a causative agent of fish mortality (Karlsen et al., 2019; John et al., 2022). Its abundance varied between 0.4 % and 51.8 % of the microeukaryotic community assessed via 18S amplicon sequencing through the HAB affected region. The highest relative abundances of focal taxon, *C. leadbeateri* – recorded in our molecular sampling campaign – were obtained in the northern portion of the HAB affected region that included Middle-Balsfjord, South-Tromsø, GSSS, and Tussøy stations (Figure 1a). The microbial community composition was relatively similar between Middle-Balsfjord, South-Tromsø, and GSSS stations with increased relative abundance of 18S ASVs and 16S ASVs indicative of dinoflagellates (class *Dinophyceae*) and order *Flavobacteriales*, respectively, towards station GSSS (Figure 1b-c). The relative abundance of ASVs belonging to the focal taxon were low in Ballangen (Ofotfjord) where the HAB was first reported to originate with high fish mortality in mid-May (Karlsen et al., 2019). The community structure from Ballangen was also distinct, compared to other samples, as it was dominated by 18S ASVs indicative of centric diatoms notably from the genus *Skeletonema* (class *Mediophyceae*) and 16S ASVs classified within the genera *Pseudoalteromonas* and *Colwellia* (order *Alteromonadales*) (Figure 1b-c and Supplementary Data S2 and S3). The relative abundance of the *C. leadbeateri* was higher in samples collected from Sagfjord stations (Anevik, Dyping, Veggfjell, and Middle-Sagfjord), south from Ofotfjord, than in the Ballangen and FSS stations (Figure 1a). The microeukaryotic community in the Sagfjord location was especially diverse, comprising multiple distinct groups of dinoflagellates and centric diatoms being the most abundant. In contrast to the other sampling locations, the 16S prokaryotic ASVs among the Sagfjord stations were predominantly comprised by the order *Pseudomonadales* (Figure 1b-c).

### Microbiome dynamics across the bloom

The microeukaryotic and prokaryotic community composition changed across the time series sampling campaign at FSS (PERMANOVA: 16S, R^2^ = 0.11 and *p* = 0.001; 18S, R^2^ = 0.19 and *p* = 0.001) and GSSS (PERMANOVA: 16S, R^2^ = 0.13 and *p* = 0.001; 18S, R^2^ = 0.13 and *p* = 0.001). These dynamic communities were also significantly dissimilar between the two time series stations (PERMANOVA: 16S, R^2^ = 0.30 and *p* = 0.001; 18S, R^2^ = 0.46 and *p* = 0.001) (Supplementary Table S3). GSSS showed higher species richness as compared to FSS as measured by the number of observed ASVs per sample day; 235–288 and 244–358 ASVs for 16S and accordingly 161–265 and 306–415 ASVs for 18S in FSS and GSSS, respectively (Supplementary Figure S6).

Among the most abundant microeukaryotes assigned to the taxonomic level class, only 18S ASVs indicative of *Cryptophyceae*, *Mediophyceae*, *Dinophyceae* and *Prymensiophyceae* (including the focal taxon) were shared between both stations (Figure 2a-b). The observed community in FSS was dominated by taxa classified as *Cryptophyceae* (9-49 % of the relative abundance), comprising mainly ASVs assigned to the genus *Teleaulax*, that showed strong temporal variability in the second half of the time series (Figure 2a and Supplementary Data S4). The predominant taxon observed from GSSS was classified as *Dinophyceae,* which accounted for 34–54 % of the relative abundance during the time series (Figure 2b). Most of the ASVs belonging to the class *Dinophyceae* were not assigned to a lower taxonomic level.

**Figure 2.**
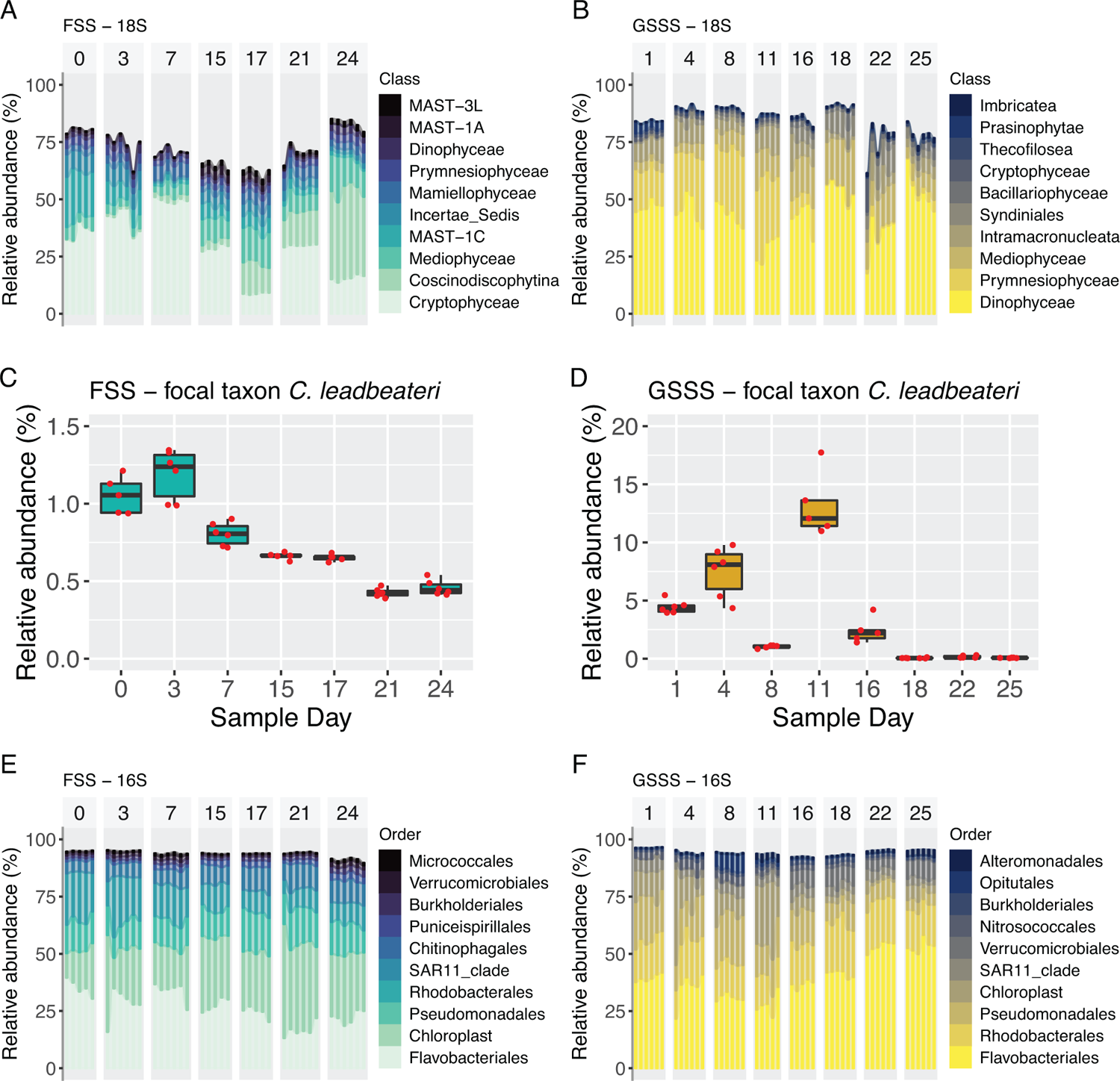
Comparative composition of the most common microeukaryotic and prokaryotic taxa collected from the time series sample stations and determined via 16S and 18S rRNA amplicon analysis. Microeukaryotes classified at the class taxonomic rank level **A**) in FSS and **B**) in GSSS. The temporal change in relative abundance of the *C. leadbeateri* focal taxon **C)** in FSS and **D)** in GSSS. Prokaryotes classified at the order taxonomic rank level **E)** in FSS and **F)** in GSSS.

A change in community structure was observed at both stations during the middle of the observed bloom; between days 7 and 15 in FSS and between days 11 and 16 at the GSSS station (Figure 2a-b). This shift was observed with an increased relative abundance of ASVs indicative of centric diatoms (class *Mediophyceae*) in both stations and the class *Coscinodiscophytina* in FSS which were almost exclusively represented by ASVs indicative within genus *Leptocylindru*s (Figure 2a-b and Supplementary Data S4). The class *Mediophyceae* comprised ASVs that were classified within different genera between stations, such as *Acrocellulus* at FSS and *Thalassiosira* at GSSS (Supplementary Data S4-S5). In FSS, the contribution of class *Prymnesiophyceae* to the community composition was minor in comparison to GSSS (Figure 2a-b) and was composed mainly by ASVs classified to the genus *Chrysochromulina* (2-3 % of relative abundance) as classified by the SILVA v138.1 database but also ASVs assigned to the *Chrysochromulina* focal taxon as verified by manual sequence alignment (0.5-1 % of relative abundance) (Figure 2c and Supplementary Data S4-S5). In GSSS, class *Prymnesiophyceae* prevailed until day 11 (42 % of the relative abundance) and thereafter drastically decreased while ASVs assigned to *Syndiniales* concurrently increased (Figure 2b). The majority of those ASVs belonging to the class *Prymnesiophyceae* were assigned to the genera *Phaeocystis* and the focal taxon (*C. leadbeateri*) (Supplementary Data S5). The pronounced decrease in abundance of the focal taxon at day 8 coincided with highest relative abundance of *Phaeocystis* (Figure 2d and Supplementary Data S5). We note here for clarity that ASVs assigned to the genus *Chrysochromulina* are not indicative of the focal taxon, even though the unialgal control has been deposited and frequently referred to as *Chrysochromulina leadbeateri* UiO-035.

The 16S ASVs – classified at the level of order – showed less temporal variability than the 18S ASVs (at the class level) across the time series in FSS and GSSS. The taxa with the highest relative abundances were the same in FSS and GSSS sample stations (Figure 2e-f). Taxa classified to the order *Flavobacteriales* showed the highest relative abundance with ASVs belonging *Rhodobacterales* and *Pseudomonadales* being common as well in both time series stations (Figure 2e-f). The FSS and GSSS locations harbored distinctly different relative abundances of genera assigned to *Flavobacteriales*. At station GSSS, the dominant genera were *Polaribacter*, *Ulvibacter*, *Formosa* and the NS5 marine group, as compared to FSS where the NS5 and NS9 marine groups were dominant (Supplementary Data S6 and S7). At GSSS, members of the order *Verrumicrobiales* increased in relative abundance (from 1 % to 8 %) within the same time frame as major changes in the microeukaryotes were observed (Figure 2f). Also, the abundance of the *Pseudomonadales* order decreased towards the end of the time series while the contribution of *Flavobacteriales* simultaneously increased (Figure 2f). The influence of Archaea to the 16S ASV composition was minor at both stations and accounted for only 0.1 % and 0.4 % of the relative abundance in FSS and GSSS, respectively (Supplementary Data S6 and S7).

Since this study did not include measurements of chlorophyll *a* to indicate changes in phytoplankton abundance, the 16S ASVs identified as Chloroplast were used as an inference. The average relative abundance of ASVs assigned to Chloroplast was nearly three times higher in FSS than in GSSS and the temporal pattern was different: the abundance increased towards the end of the time series in FSS whereas in GSSS this was opposite (Figure 2e-f).

### Environmental variability during the HAB

The surface water properties – temperature and salinity – were relatively similar during the HAB, as observed from both time series stations (FSS and GSSS) at 10 m depth from where microbial biomass was collected (Supplementary Table S4). The overlaying surface water was more stratified in FSS than GSSS. This was due to a low salinity layer (< 30) which persisted near the surface during the entire time series simultaneously with a strong increase in observed temperature (Supplementary Figure S1 and S2). The mixed layer depth was observed near the surface (3 m) throughout the study period in FSS and in the beginning in GSSS time series, where it then sank down to 7 m by the end of the GSSS time series while the surface water became more homogeneous after day 8 (Supplementary Figure S1 and S2). The mean NO_3_^-^ + NO_2_^-^ concentration was significantly higher in GSSS than in FSS (0.43 and 0.18 µmol L^-1^, respectively; Tukey, p < 0.02) whereas the Si(OH)_4_ and PO_4_^3-^ concentrations did not show statistically significant differences between stations (Supplementary Table S4).

### Beta diversity in context with the environment

Both the microeukarytotic and prokarytotic components of the HAB-associated microbiome were dissimilar between GSSS and FSS stations and throughout the observed time series. This was measured via unweighted UniFrac distances that were ordinated by dbRDA analysis to infer the correlation between selected environmental measurements to differences in community composition (Figure 3). Results of beta diversity also corroborated the observed change in community structure during the middle of the time series that was seen and noted above by inspecting changes in relative taxonomic abundance (Figure 2). The dbRDA analysis (i.e., sum of RDA1 and RDA2) showed that 77.9% and 83.7 % of the total variation in the prokaryotic and microeukaryotic community composition was correlated to environmental measurements during the time series (Figure 3). Most of this variation was explained by the first RDA component (RDA1: 16S = 60.8 % and 18S = 72.6 %) inferring that the variation of both prokaryotic and microeukaryotic communities between FSS and GSSS may be driven by response to environmental factors such as MLD, salinity, and NO_3_^-^ + NO_2_^-^ (Figure 3). Whereas the temporal variability within stations was mainly explained by the second RDA component (RDA2) which accounted for 17.1 % and 11.1 % of the variance in prokaryotic and microeukaryotic communities, respectively.

**Figure 3.**
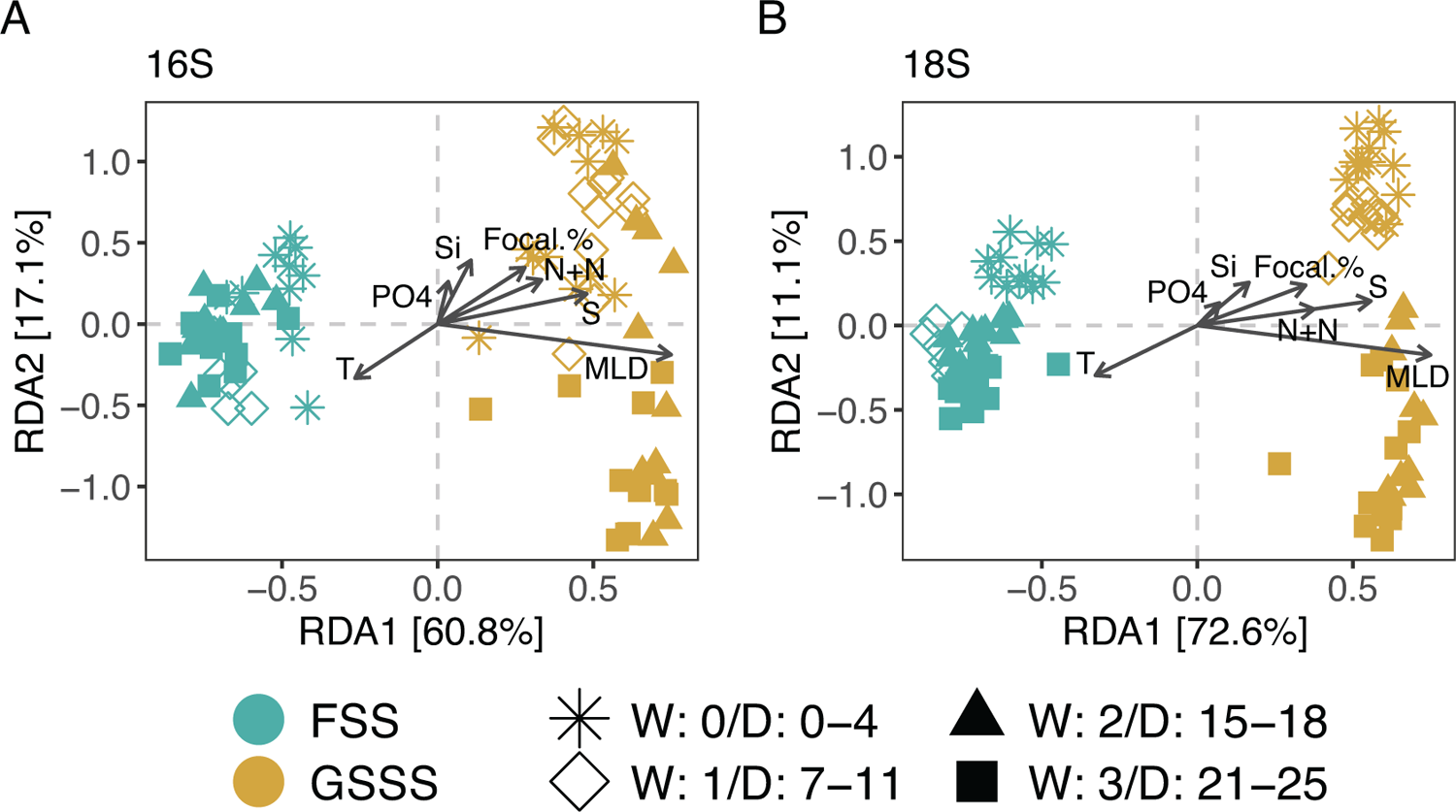
Dissimilarity in prokaryotic and microeukaryotic community composition between sample stations was inferred via beta-diversity as measured with unweighted UniFrac distances and related to environmental measurements. Distance-based redundancy analysis (dbRDA) biplot of **A)** prokaryotes and **B)** microeukaryotes. Color represents station and shape comprises samples (sample scores) within each sample week according to legend information. Arrows indicate environmental factors and point to the direction of maximum variation of the respective factor. T, temperature; S, salinity; MLD, mixed layer depth, Si, Si(OH)_4_; PO4, PO_4_^3-;^ N+N, NO_2_^-^ + NO_3_^-^; Focal %, relative abundance of the *C. leadbeateri* focal taxon in 18S data set.

### Temporal trends and co-blooming taxa

The marine microbiome at the two time series stations, FSS and GSSS, was complex and consisted of a minimum of 235 and 161 16S and 18S ASVs, respectively, each maintaining its own dynamic trajectory during the observed time series. Examination and subsequent clustering of the most predominant blooming patterns was performed via the k-medoids clustering method. The resulted number of clusters per data set and time series station, as assessed via the Calinski-Harabasz index (normalized ratio for inter-intra-cluster variance) (Supplementary Figure S5), was 3, except for 18S at FSS where 4 clusters were selected. The temporal pattern of each cluster was specified by its medoid ASV as a representative shape for cluster’s blooming dynamics of microeukaryotes (Figure 4a-b) and co-blooming taxa assigned to 16S ASVs (Figure 5a-b).

**Figure 4.**
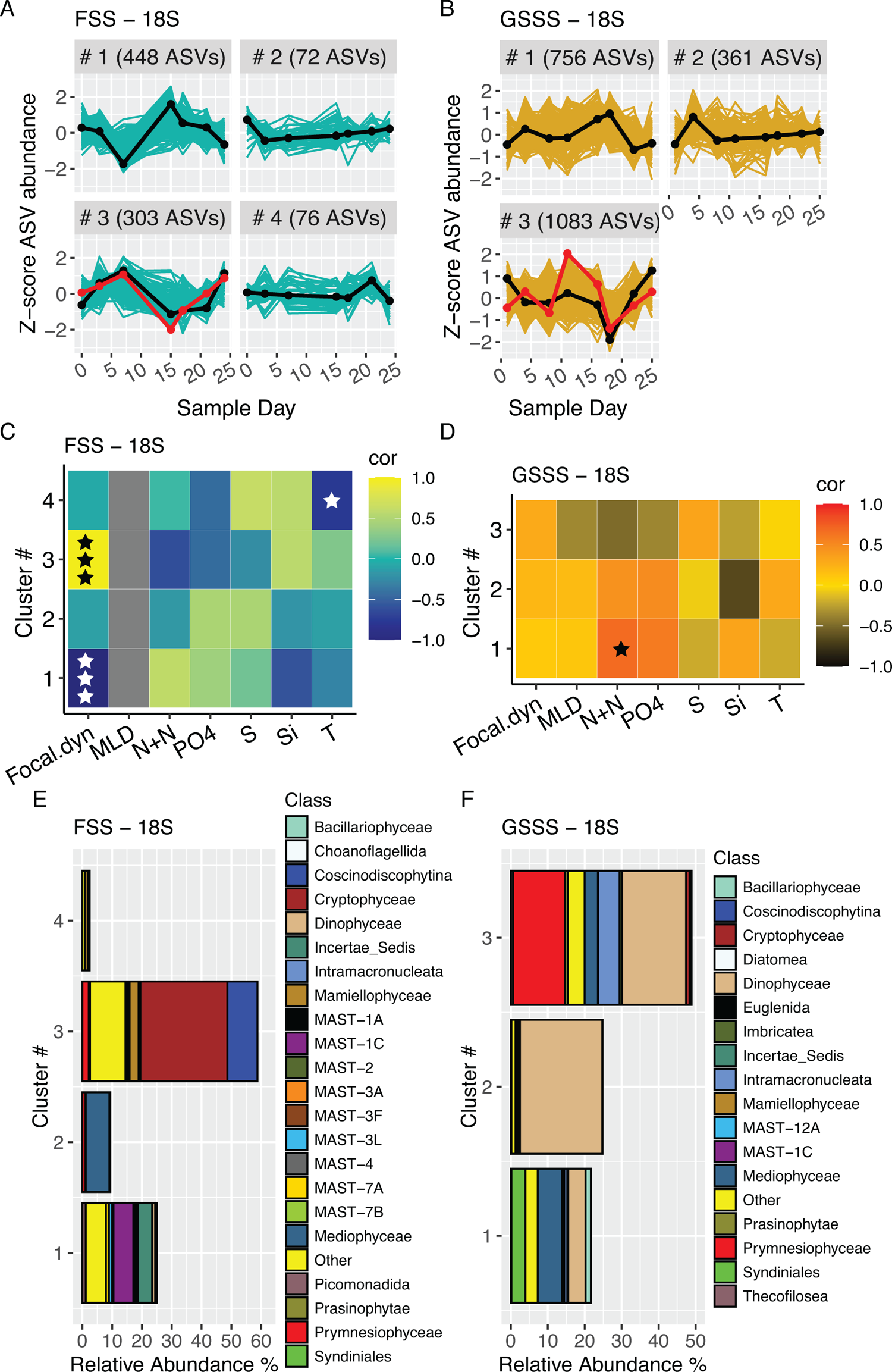
Characterization of major blooming patterns in microeukaryotic communities assessed by k-medoid clustering analysis with z-transformation and detrending of 18S ASVs. Temporal dynamics of microeukaryotic taxa and number of ASVs per defined cluster **A)** in FSS and **B)** in GSSS station. The blooming pattern of each cluster is specified by medoid taxon and drawn with black line. The red line specifies the blooming pattern of focal ASV. Y-axis is a z-score and a value 0 denotes the mean abundance. Heatmap of Pearson’s correlation between temporal dynamics of medoid taxon (z-scores) of each cluster and focal ASV (z-scores) and environmental factors **C)** in FSS and **D)** in GSSS. Color indicates Pearson’s correlation coefficient according to the color legend and level of significance is marked with ★ (★, *p* ≤ 0.1; ★★, *p* ≤ 0.05; ★ ★ ★, *p* ≤ 0.001). The grey color for MLD in FSS denotes undefined correlation as MLD values remained the same through the study period. Focal.dyn, temporal dynamic of focal ASV; MLD, mixed layer depth; N+N, NO_2_^-^ + NO_3_^-^; PO4, PO_4_^3-^;S, salinity; Si, Si(OH)_4_; T, temperature. Taxonomic profile of the most common microeukaryotes of each cluster as relative abundance in a whole community classified at the class taxonomic rank level **E)** in FSS **F)** in GSSS.

**Figure 5.**
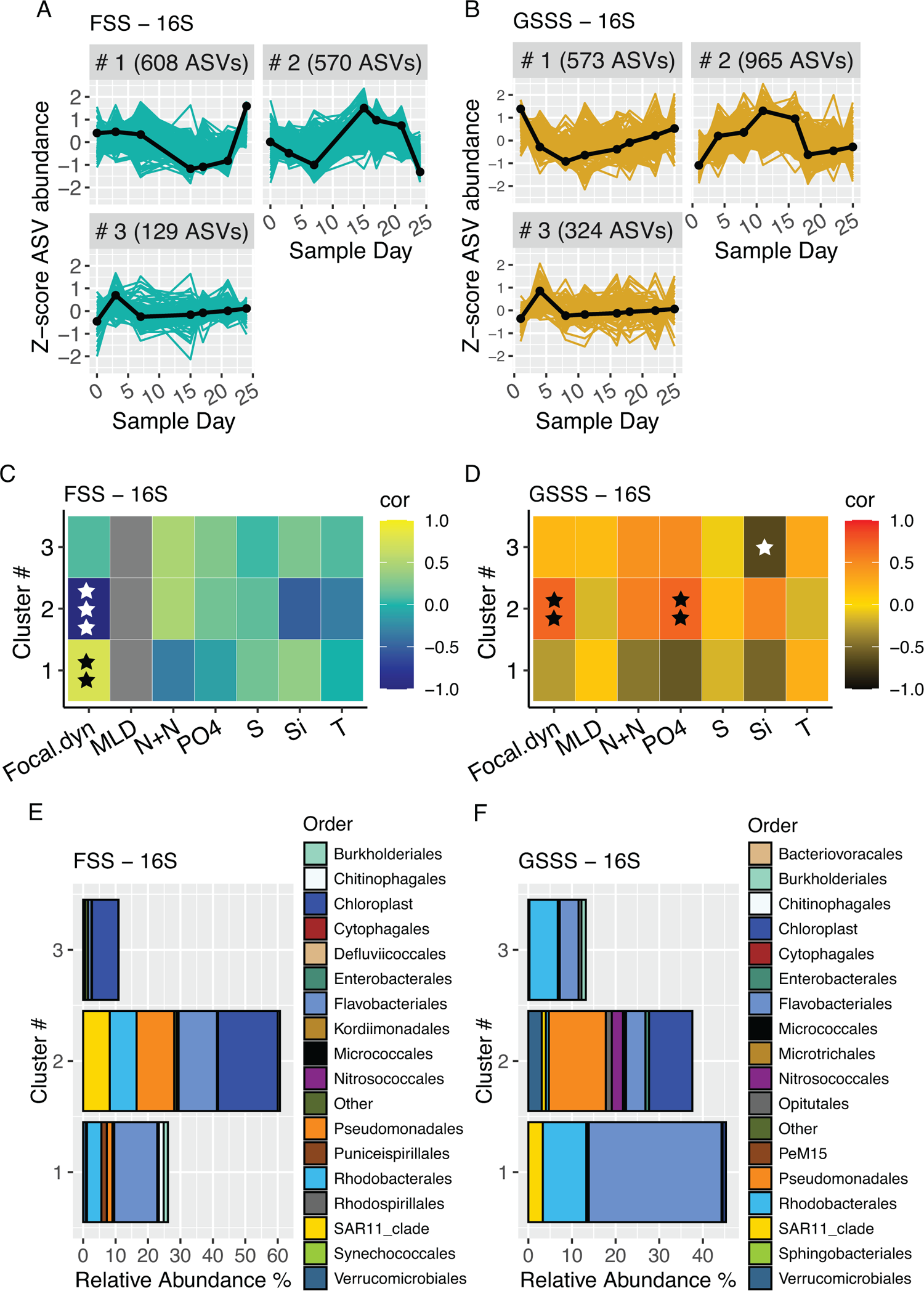
Characterization of major blooming patterns in prokaryotic communities assessed by K-medoid clustering analysis with z-transformation and detrending of 16S ASVs. Temporal dynamics of microeukaryotic taxa and number of ASVs per defined cluster **A)** in FSS and **B)** in GSSS station. The blooming pattern of each cluster is specified by medoid taxon and drawn with black line. Y-axis is a z-score and a value 0 denotes the mean abundance. Heatmap of Pearson’s correlation between temporal dynamics of medoid taxon (z-scores) of each cluster and focal ASV (z-scores; Figure 4) and environmental factors **C)** in FSS and **D)** in GSSS. Color indicates Pearson’s correlation coefficient according to the color legend and level of significance is marked with ★ (★, *p* ≤ 0.1; ★★, *p* ≤ 0.05; ★ ★ ★, *p* ≤ 0.001). The grey color for MLD in FSS denotes undefined correlation as MLD values remained the same through the study period. Focal.dyn, temporal dynamic of focal ASV; MLD, mixed layer depth; N+N, NO_2_^-^+ NO_3_^-^; PO4, PO_4_^3-^; S, salinity; Si, Si(OH)_4_; T, temperature. Taxonomic profile of the most common prokaryotes of each cluster as relative abundance in a whole community classified at the class taxonomic rank level **E)** in FSS **F)** in GSSS.

The medoid 18S ASV of cluster 1 and 3, from both stations, exhibited approximately inverse blooming patterns. These clusters also showed strong fluctuation in relative abundances across the time series, including a strong positive and negative shift, respectively, around the middle time points and negative correlations to environmental measurements (Figure 4a-d). The medoid taxon of cluster 2 and 4 in FSS and cluster 2 in GSSS exhibited a short increase in abundance in the beginning or end of time series (Figure 4a-b).

The focal ASV was included within cluster 3, at both stations (Figure 4a-b). There were several microalgae that co-bloomed with the focal ASV, but these were distinct between the FSS and GSSS stations (Figure 4). In FSS the focal ASV co-bloomed with genera *Teleaulax* (class *Cryptophyceae*) and *Leptocylindrus* (class *Coscinodiscophytina*) which comprised 16-53% and 1-53 % of the relative abundance within their respective cluster 3 (Figure 4e and Supplementary Data S8). A diverse composition of ASVs indicative of MAST clades (marine stramenopiles) were also detected, although their relative abundances were minor (< 1 % of relative abundance within the cluster 3). This group was represented with other nano- and picoplankton such as genera *Telonema* (class *Insertea Sedis*) and *Picomonas* (class *Picomonadida*) the taxa in cluster 1 with inverse blooming pattern to cluster 3 (Figure 4c, e and Supplementary Data S9). ASVs from the genus *Phaeocystis* predominantly co-bloomed at GSSS with the focal ASV (within cluster 3) together with a less abundant group of ciliates (class *Intramacronucleata*) and members belonging to genus *Thalassiosira* (class *Mediophyceae*) (Figure 4d, f and Supplementary Data S10). ASVs belonging to the class *Dinophyceae* also included in cluster 3 from GSSS (along with the focal ASV), and these were composed mainly of members classified within the genus indicative of *Gymnodinium*.

The blooming patterns of 16S ASVs were also clustered. Among these, two clusters (1 and 2 from FSS and GSSS, respectively) showed significant positive correlation (FSS: *r* = 0.79, *p* = 0.036 and GSSS: *r* = 0.71, *p* = 0.047) with the focal ASV. Only cluster 2 from station FSS showed a significant negative correlation with the focal ASV (*r* = −0.96, *p* < 0.001) (Figure 5c-d). The temporal trend represented by the medoid ASV was nearly identical in cluster 3 at both stations corresponding to the blooming pattern of cluster 2 and inverse blooming pattern of cluster 4 for microeukaryotes in GSSS and FSS, respectively.

We infer that the bacterial taxa from the GSSS station belonging to cluster 2 co-bloomed with phototrophic protists in general, because the ASVs assigned to Chloroplasts occupied the same cluster (Figure 5f and Figure 4f). This cluster was also the only one to show a statistically significant correlation with phosphate concentrations. The same pattern was not observed in FSS where approximately two thirds of Chloroplast ASVs were identified within cluster 2 and rest in cluster 3 (Figure 5e).

Pearson correlation enabled us to infer that different bacterial communities co-bloomed with *C. leadbeateri* focal taxon, corresponding to the observations within 18S microeukaryotes (Figure 5c-d). Cluster 1 at station FSS was dominated by the order *Flavobacteriales* (35-63 % relative abundance within cluster 1) (Figure 5e) and was predominated by sequences identified within the NS5 marine group but also *Ulvibacter* (Supplementary Data S11). ASVs assigned to *Rhodobacterales*, notably evenly dominating genera *Amylibacter* and *Planktomarina*, also shared the similar temporal dynamics with the focal ASV (Figure 5e and Supplementary Data S11). The positive co-blooming trend between *Flavobacteriales* and *Rhodobacterales* classes and the focal ASV was not observed at station GSSS.

## DISCUSSION

The results from this study show distinct spatial variation and differences in the dynamic microbiome community composition across the portion of the northern Norwegian coast that was strongly affected during the 2019 *Chrysochromulina leadbeateri* HAB. This was a major and disruptive event due to severe fish mortality and hence there were several sampling campaigns launched simultaneously by public and private stakeholders. This study provides a molecular-based investigation of the holistic microbiome during the 2019 HAB event and sets the blooming pattern of *C. leadbeateri* within the context of its co-occurring community members.

### Indication for separate focal blooms

The 2019 HAB was reported on and monitored daily with focus on afflicted fish farm localities by the Norwegian Directorate of Fisheries (Karlsen et al., 2019). The results obtained from our amplicon-based sequencing approach are expected to be more complex and distinct from the concurrent morphology-based taxonomy and cell counting surveys. However, our results share some reassuring similarities to the reported microcopy cell counts but also 18S amplicon sequence reads presented in a recent study indicating that high *C. leadbeateri* focal taxon abundance was found in Balsfjord (Karlsen et al., 2019; John et al., 2022). The current study did not capture the onset of the HAB due to a reactive sampling effort to an event that is (at least today) impossible to predict.

According to the prevailing consensus based on reported fish mortality, the toxic bloom of the focal taxon initiated in Ofotfjord. The last report on fish mortality in Ofotfjord was dated four days before the sample in Ballangen was taken (Karlsen et al., 2019), thus there was, presumably, a mismatch between the bloom peak and time of sampling. We do not have data that represents the community composition during the first reports of fish mortality. This may explain why we found a surprising high disparity between expected (i.e., high focal taxon abundance) and observed microbial community composition at Ballangen, which did not correspond with any other sampled microbial community composition (Figure 1). The microeukaryotic community was dominated by diatoms, nearly exclusively composed by *Skeletonema*. A similar shift from *C*. *leadbeateri* presence to absence with concurrent predominance of diatoms was observed during the 1991 HAB around Ofotfjord and surrounding areas (Hegseth and Eilertsen, 1991; Rey and Aure, 1991). Correspondingly, the sampled prokaryotic community composition in Ballangen does not presumably represent either the situation with *C. leadbeateri* as a disruptive causative agent as several genera belonging to the observed predominant bacterial order *Alteromonadales* have been found to associate with *Skeletonema* sp. (Deng et al., 2021).

The focal taxon *C. leadbeateri* is a common and natural member of phytoplankton community along the entire northern Norwegian coast, although its spatial and interannual variation has not been well described, as only two following years of monitoring was conducted in area south and north from Ofotfjord after the 1991 HAB (Hegseth and Eilertsen, 1991; Heidal and Mohus, 1995). The present and recent study, however, provides evidence for high spatial variation and presumably non-cell-density dependent toxicity (John et al., 2022). As mentioned, the high abundance of *C. leadbeateri* in the Balsfjord area was corroborated by manual cell counts and revealed by 18S rRNA amplicon sequencing of this study two weeks after the first reported fish mortalities from Ofotfjord (Figure 1a) (Karlsen et al., 2019). Thus, this raises the question whether a bloom of focal taxon commenced locally in Balsfjord. Due to limited connectivity, it is likely that *C. leadbeateri* blooms developed independently in both fjords since its relative abundance was low in FSS station (Figure 2c) and no afflicted fish farms were reported near the FSS station, despite of the affect on fish farms in the area north of Ofotfjord (Karlsen et al., 2019). Interestingly, despite high abundance of the focal taxon and effective advection-driven dispersal of cells from the inner part of Balsfjord towards GSSS and Tussøya stations, only one fish farm near Tussøya was afflicted (Figure 1a). However, this farm experienced limited impacts, and its relation to the HAB remained somewhat questionable (Karlsen et al., 2019). This observation also lends some indications to the idea that there may have been at least two distinct physiological states between these two *C. leadbeateri* blooms, and that only one of them was “harmful”.

### Localized microbial communities

One of the major questions that drove this study was to ask how the marine microbiome composition compared across locations during the HAB time series. The results revealed distinct communities between sample locations, each possessing different predominating bacterial and eukaryotic microbial groups (Figure 1 and Figure 2). This observation on spatial heterogeneity provides new information on the development of the late bloom phase within an interconnected coastal area (Wassmann et al., 1996). GSSS was the only station where *Phaeocystis* abundance exceeded that of the focal taxon. In fact, the relative abundance of *Phaeocystis* was less than 1 % at all other stations, which was an unexpected result given previously reported ubiquity and prolonged prevalence (March-August) in this region (Eilertsen et al., 1981; Degerlund and Eilertsen, 2010).

Although the predominant microalgae groups found in the two time series stations were different (FSS: cryptophytes, *Leptocylindrus*, heterotrophic nanoflagellates; GSSS; dinoflagellates, prymnesiophyte, *Thalassiosira*), the most abundant bacterial taxa – i.e. orders *Flavobacteriales*, *Rhodobacterales* and *Pseudomonadales* – were essentially consistent. Previously reported studies have also shown that members belonging to these orders commonly accompany microalgae in marine environments, except in the case of *Teleaulax* sp. and *C. leadbeateri,* which may be due to lack of previous records from events such as the 1991 HAB of northern Norway (Teeling et al., 2016; Ajani et al., 2018; Zhou et al., 2018; Aalto et al., 2022). The bacterial community compositions obtained from Middle-Balsfjord and South-Tromsø stations where distinct and corresponded to the highest observed relative abundance of the focal taxon. Specifically, they harboured high relative abundances of SAR11 (mainly clade Ia) and low contributions of *Flavobacteriales,* which is interesting given that SAR11 tend to be negatively correlated with copiotrophic bacterial groups before and after the primary phytoplankton bloom (Needham and Fuhrman, 2016; Teeling et al., 2016). However, the single time point stations present a major limitation for interpreting the HAB microbiome, given that the time series stations (FSS and GSSS) showed pronounced temporal variability during the event.

### Temporal community dynamics

Another major aim of this study was to identify the temporal patterns of the HAB associated microbiome and attempt to determine which taxa co-bloomed with the focal ASV. We found that most of the microeukaryotic and bacterial taxa either co-bloomed or inversely co-bloomed with the focal ASV – i.e., the clusters that showed significant positive or negative correlation with the focal ASV also had the highest taxonomic richness (Figure 4 and Figure 5). There has been evidence that the most abundant taxa undergo stronger shifts in their abundance than rare taxa, which corresponds with our results where the most abundant bacterial orders and microeukaryotic classes were mainly found within the clusters of highest temporal variability (Lindh et al., 2015). However, the high number of taxa in these clusters is somewhat surprising with regard of previous findings of highly coherent communities with well-defined groups of organisms with the potential to interact (Martin-Platero et al., 2018). It is likely that the prescribed temporal (sampling) resolution of this study was too coarse to capture the true frequency of taxonomic turnover, which can be very fast (Needham and Fuhrman, 2016; Martin-Platero et al., 2018). However, when an unpredicted and regionally large (250 km wide) HAB occurs, it poses major logistical challenges to react quickly and access the sample locations with frequency that would be the most optimal with biological perspective. The choice of optimal sampling locations is also limited by immediate accessibility to sampling resources, such as suitable watercraft.

### The environment

The third major question of this study was related to the abiotic environment and how it may have influenced the microbiome composition and presence of the *C. leadbeateri* focal taxon. The HAB – as inferred from relative abundance of the focal taxon – was stronger at GSSS than at FSS, yet it occurred concurrently at both stations. The strong decrease in *C. leadbeateri* abundance in the middle of time series coincided, especially at GSSS, with a change in community composition. However, this was evidenced poorly by environmental factors at both stations. Intriguingly, these results are in accordance with previous findings suggesting lower importance of continuous environmental factors such as temperature, inorganic nutrients, and chlorophyll *a* on a change in community composition after seasonal bloom initiation than biotic interactions (Lindh et al., 2015; Needham and Fuhrman, 2016). Whereas, a strong periodical physical forcing (wind, turbulence) can cause a change of entire community (Martin-Platero et al., 2018). The measured water column properties did not indicate strong vertical mixing. However, the surface water became somewhat more homogenous after the first week in the time series at GSSS as revealed from CTD-profiles (Supplementary Figure S2).

The difference in microeukaryotic composition between the two time series stations may be related to environmental factors as revealed by dbRDA analysis (Figure 3). The phosphate concentration was low at both stations and in addition the NO_3_^-^ + NO_2_^-^ level was low in FSS, which potentially promoted the development of communities of heterotrophic and mixotrophic protists. Several MAST-clades (e.g., MAST-1C as dominant in FSS) are known to be bacterivores and *Teleaulax* has been determined to be mixotrophic, feeding heterotrophically on bacteria, thus relaxing the need for inorganic nutrients (Massana et al., 2006; Du Yoo et al., 2017). It is difficult to draw a link between nutrients and the focal taxon with the current data due to lack of nutrient measurements from stations with high focal taxon abundance such as Middle-Balsfjord and South-Tromsø. Despite the nutrient information, a plausible connection would be unlikely as even a 14-year survey in southern Norway on *Chrysochromulina* spp. cell and nutrients concentration did not provide a clear connection (Dahl et al., 2005). However, there are indications that *Chrysochromulina* spp. thrive in high N:P ratios during low phosphate concentration which potentially promotes toxicity if the condition becomes phosphate limited (Dahl et al., 2005; Edvardsen and Imai, 2006).

## CONCLUSION

A summary of investigations of the 1991 *Chrysochromulina leadbeateri* related harmful algal bloom in northern Norway writes that it would not have been possible to predict such an incident but it would be necessary to launch a coastal monitoring program (Rey, 1991). Three decades later, we fully agree with the latter statement. We certainly have a long way to go in order to predict how ecological principles underpin this sporadic event in a highly variable coastal area and it is difficult to get there by performing investigations that simply react to only those blooms that have acute economic impacts. It is also difficult to plan to observe such a bloom that happens so infrequently. It has become highly evident that this type of reoccurring bloom is extremely destructive because it causes substantial losses for the fish farmers. In 1991 the fish kill damage linked to *C. leadbeateri* was 742 tonnes with a corresponding direct short term economic cost of 3.5 million USD, while in 2019 these values were increased to 14 500 tonnes and more than 100 million USD (Karlson et al., 2021). Thus, it is expected these numbers will become even higher in the future with increasing aquaculture industry, and there is no reason to believe that this type of HAB will not reoccur. It is challenging to draw any conclusions about exact causation behind the 2019 HAB, especially since the natural bloom cycles of *C. leadbeateri* are not well understood. Therefore, this study relying on molecular-based methods provides a first insight on the spatial and temporal variability of the *C. leadbeateri* focal taxon together with its associated marine microbiome. It also supports future molecular-based studies by providing a genetic link with currently available taxonomic annotation between current and year 1991 *C. leadbeateri*. Otherwise, the *C. leadbeateri* focal taxon may be overlooked due to database limitations – an issue also present in this study. We found that most of the taxa – including the most abundant members of prokaryotes and microeukaryotes and the focal taxon – underwent strong and rapid variability in their succession dynamics. These temporal dynamics were poorly connected to the environmental conditions measured, suggesting that other factors such as biological interactions might drive the late bloom dynamics.

## AUTHOR CONTRIBUTIONS

NJA and HCB conceptualized the study. NJA, EG-M, JBS, LD, CJH and RI carried out the sampling. NJA, HS, SK and SP performed lab work and processed amplicon sequence analyses. NJA and HCB conducted data analysis and NJA wrote the manuscript draft. All authors contributed to the final version.

## ACKNOWLEDGMENTS

We greatly appreciate the logistical support by staff at Lerøy Aurorás aquaculture facility at Solheim. We also acknowledge the collaboration between Finnfjord AS and UiT-The Arctic University of Norway for lending sampling infrastructure and support. We would like thank Åge Mohus and Vigdis Tverberg at Nord University for their help with sample coordination in southern portion of the HAB affected area. We kindly acknowledge assistance from the Environmental Sample Preparation and Sequencing Facility in the Biosciences Division of the Argonne National Laboratory.

## SUPPLEMENTARY MATERIAL

**Supplementary Figure S1.**
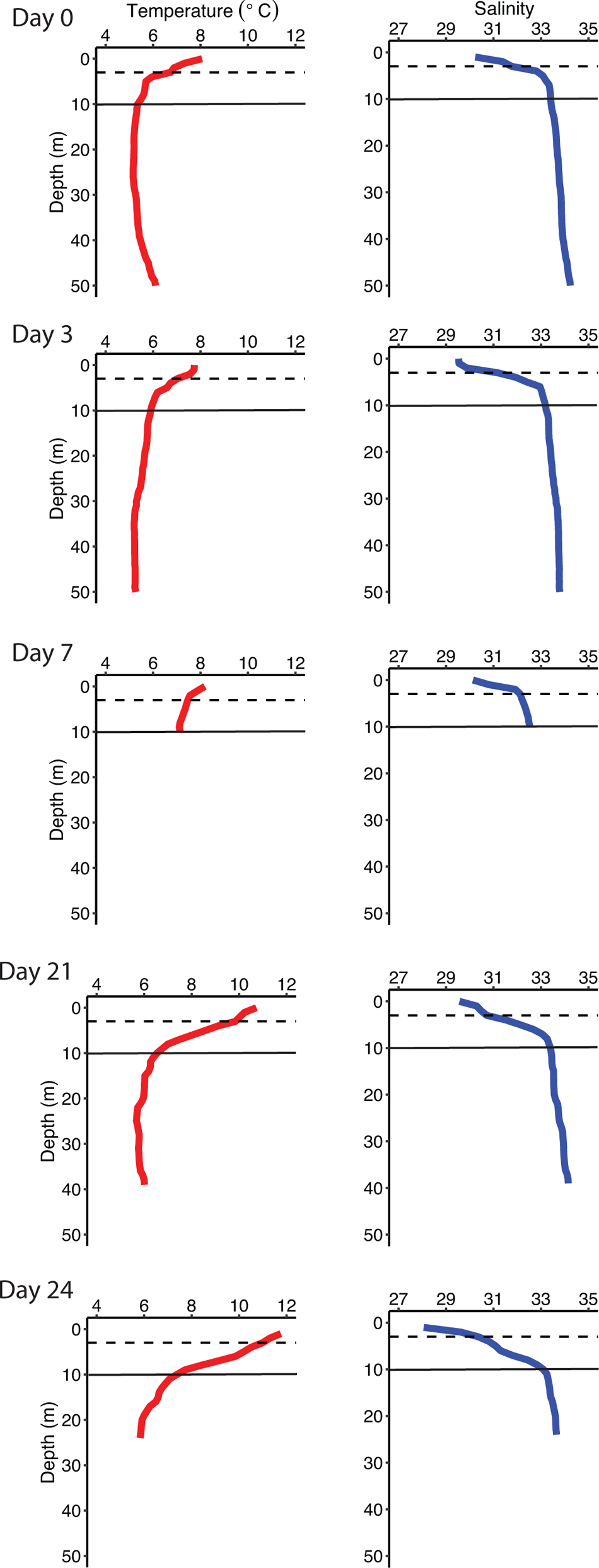
Surface water profiles of temperature and salinity obtained from CTD-casts in FSS time series station. The dashed line denotes calculated mixed layer depth (MLD) based on density gradient and solid line marks the depth of sampled microbial communities (10 m). Note missing CTD-casts in sample days of 15 and 17.

**Supplementary Figure S2.**
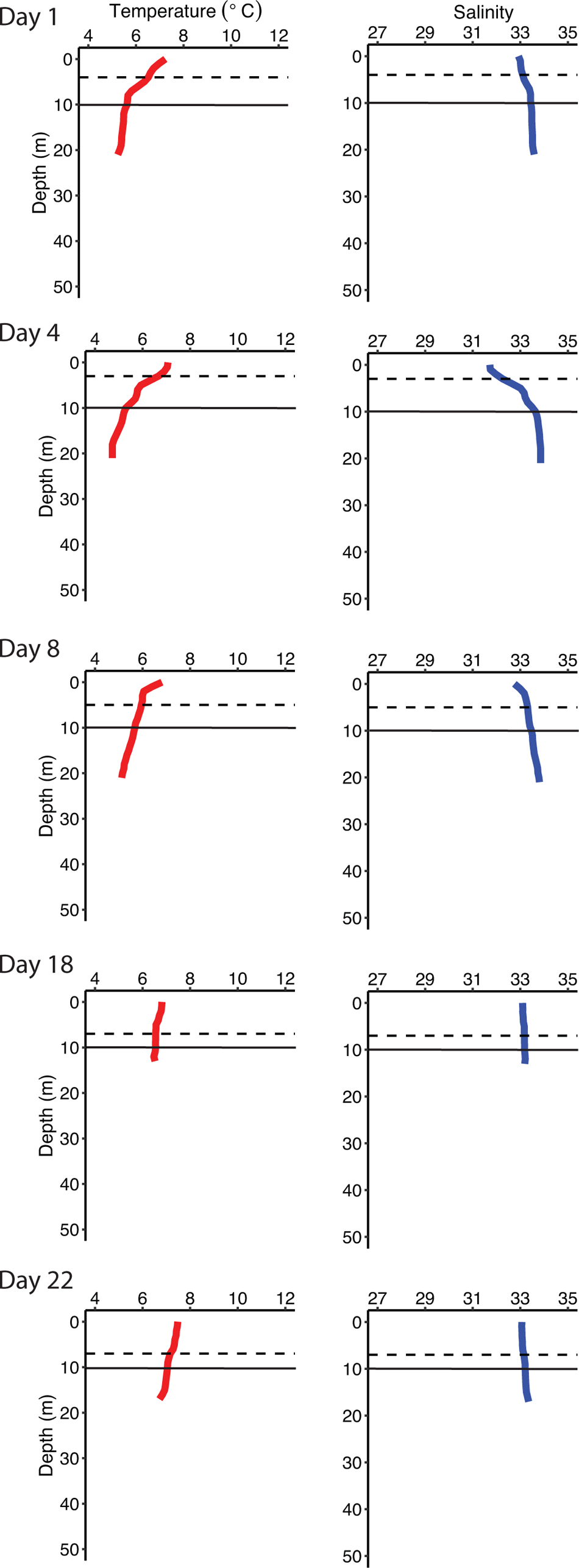
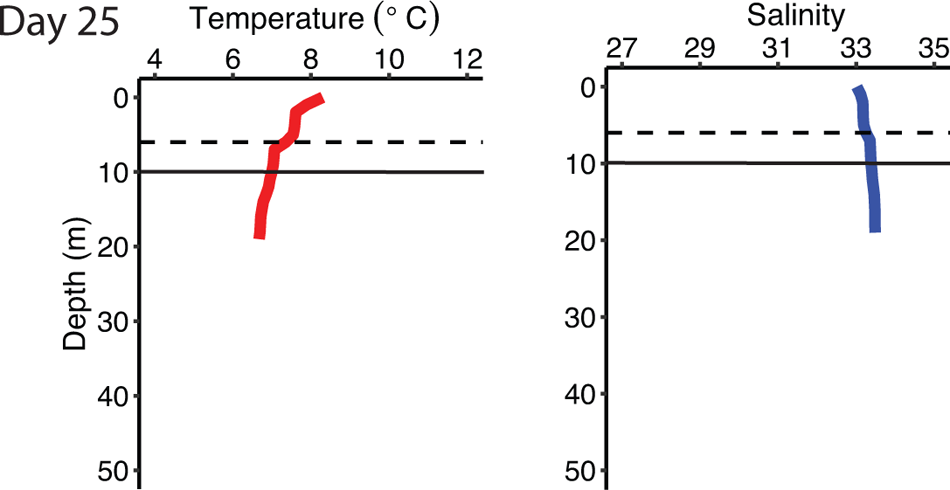
Surface water profiles of temperature and salinity obtained from CTD-casts in GSSS time series station. The dashed line denotes calculated mixed layer depth (MLD) based on density gradient and solid line marks the depth of sampled microbial communities (10 m). Note missing CTD-casts in sample days of 11 and 16.

**Supplementary Figure S3.**
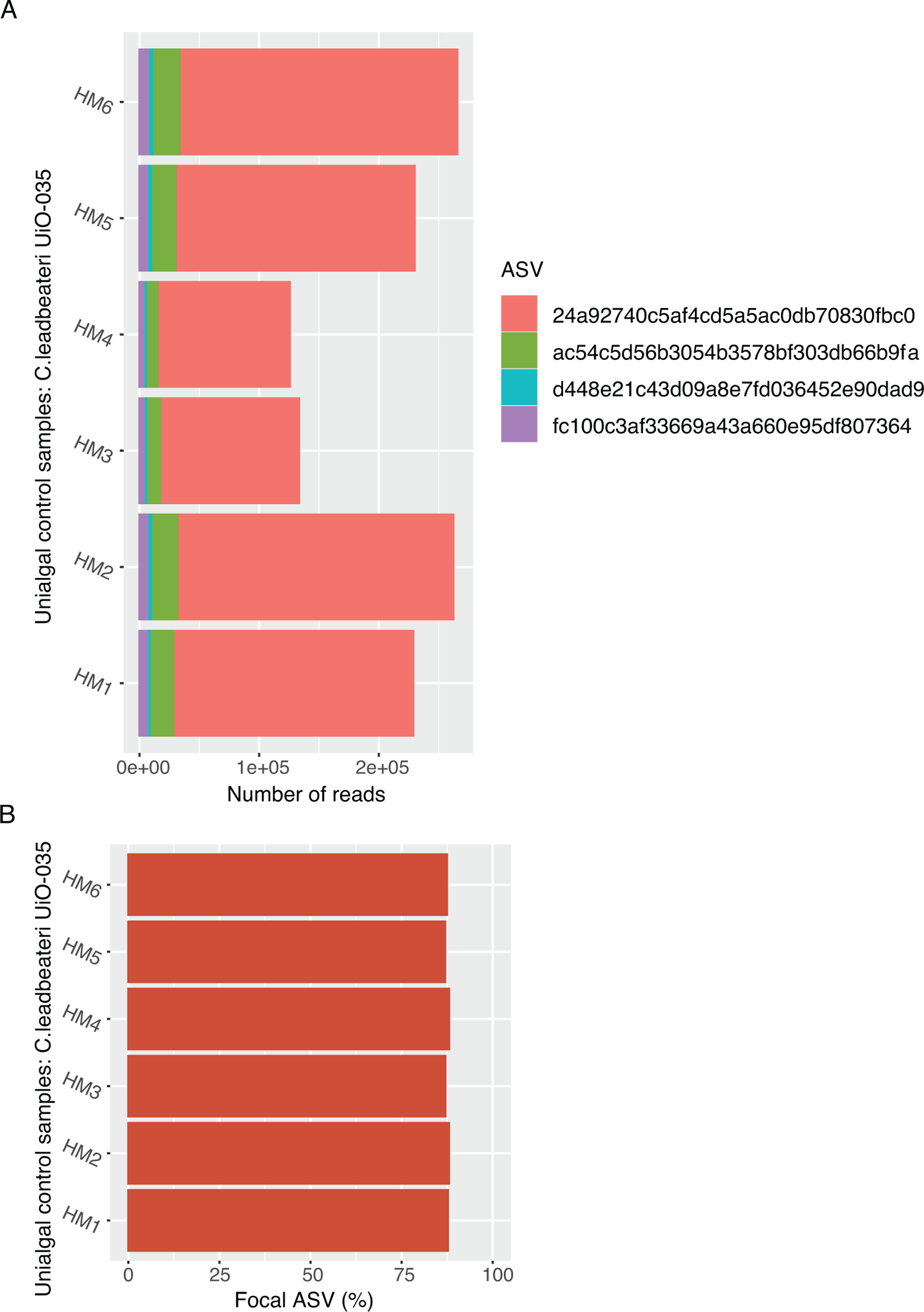
A) Number of amplicon sequence reads of each ASV assigned to OLI16029, in unialgal *Chrysochromulina leadbeateri* UiO-035 control culture. The most abundant ASV: *24a92740c5af4cd5a5ac0db70830fbc0* is referred as the “focal ASV”. B) Relative abundance of focal ASV among ASVs shown in A).

**Supplementary Figure S4.**
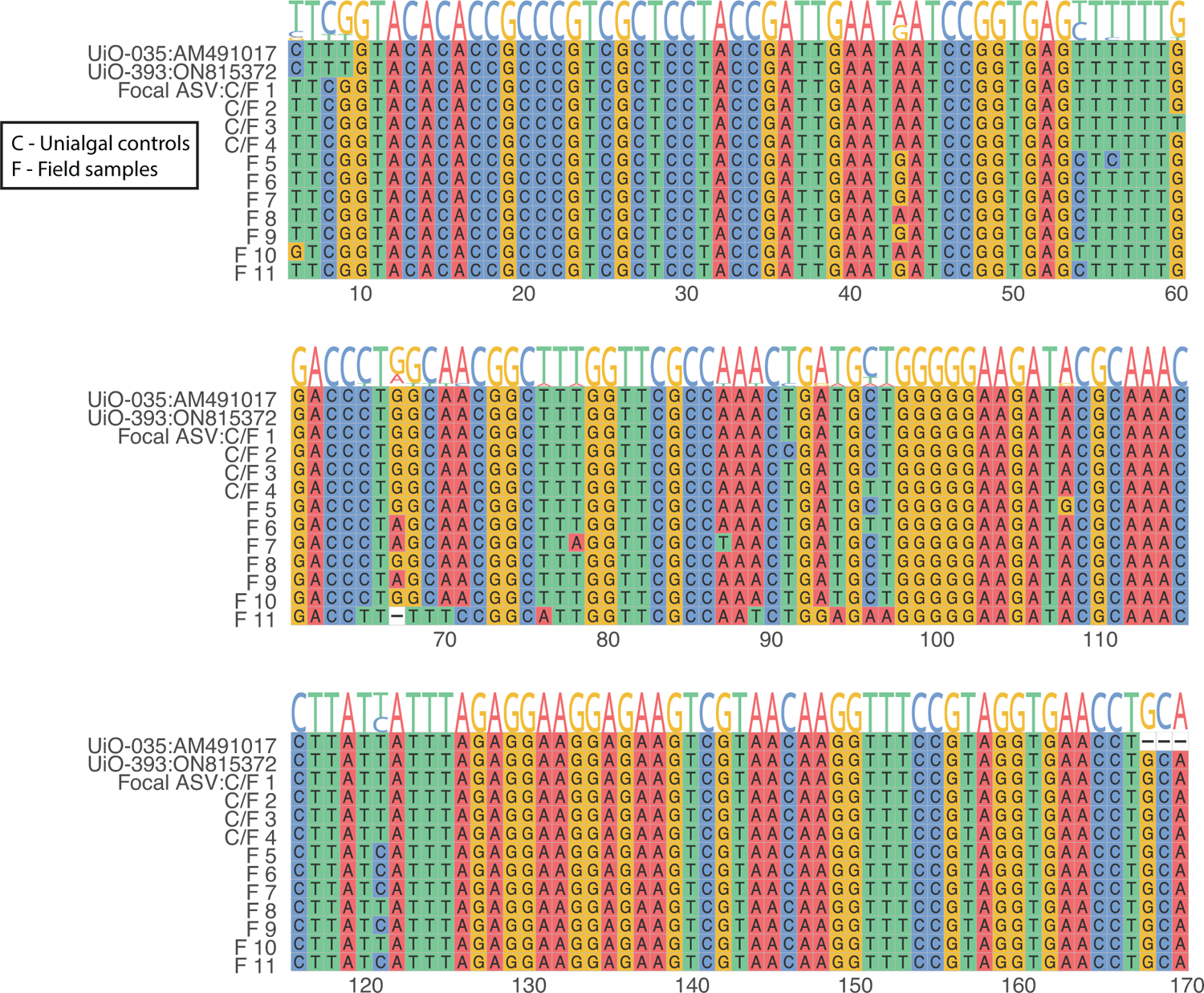
18S rRNA gene sequence alignments (MAFFT v7, strategy L-INS-i) between *Chrysochromulina leadbeateri* strains of UiO-035 (isolated from 1991 HAB) and UiO-939 (isolated from 2019 HAB) and ASVs of this study both from unialgal control cultures and field samples assigned as OLI16029 by SILVA v138.1. The sequences of strains UiO-035 and UiO-939 were obtained from Genbank under accession number AM491017 and ON815372, respectively. ASVs present in unialgal control cultures are denoted with C and in field samples denoted with F. **Focal ASV:C/F 1**: ASV *24a92740c5af4cd5a5ac0db70830fbc0*; **C/F 2**: ASV *ac54c5d56b3054b3578bf303db66b9fa*; **C/F 3**: ASV *d448e21c43d09a8e7fd036452e90dad9*; **C/F 4**: ASV *fc100c3af33669a43a660e95df807364*; **F 5**: ASV *2c35c44a2fccfc37189c2069cbe1a055*; **F 6**: ASV *339b61ffec8c56a7432a1e8e4c4b38aa*; **F 7**: ASV *4f7b870b345b622629b6aaa8a8ad5496*; **F 8**: ASV *95d93ff25078e3eccfbc0bdf68635ed0*; **F 9**: ASV *d0099e62abb8d798ee68455fe7823601*; **F 10**: ASV *daa2cdc1dcfb31d58216538504d7fe8a*; **F 11**: ASV *fd29fa001773f2999f28b9012e132556*. Note that the sequences are continuous although shown as stacked.

**Supplementary Figure S5.**
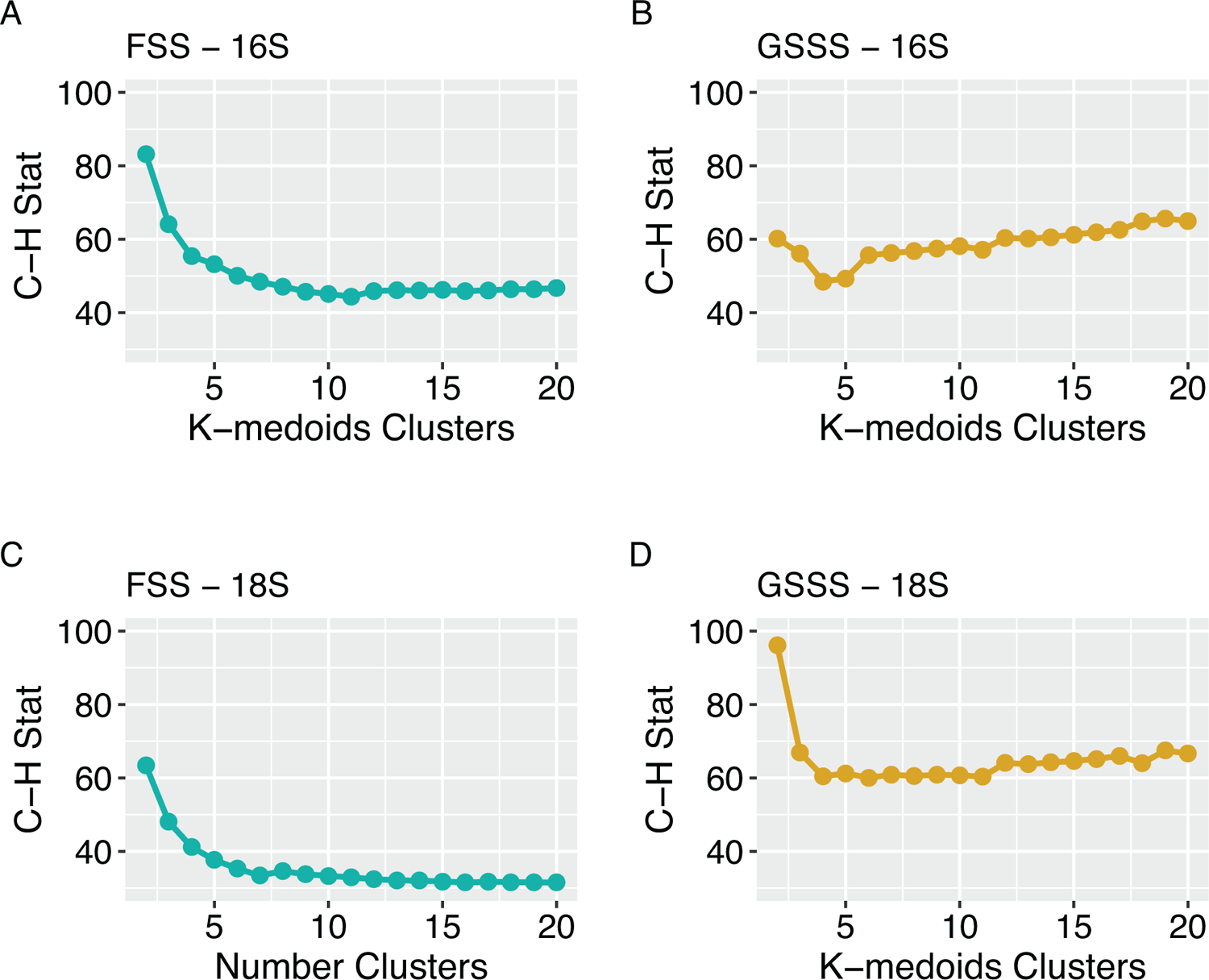
The Calinski-Harabasz index (y-axis, C-H Stat) was used to determine the number of K-medoids clusters. C-H Stat is a normalized ratio of the inter and intra cluster variance calculated for each sample-specific K-medoids cluster. High C-H Stat value indicates more uniquely portioned clusters. A) Three clusters were chosen to represent 16S ASVs from FSS time series. B) Three clusters were chosen to represent 16S ASVs from the from GSSS time series. C) Four clusters were chosen to represent 18S ASVs from FSS time series. D) Three clusters were chosen to represent 18S ASVs from the from GSSS time series.

**Supplementary Figure S6.**
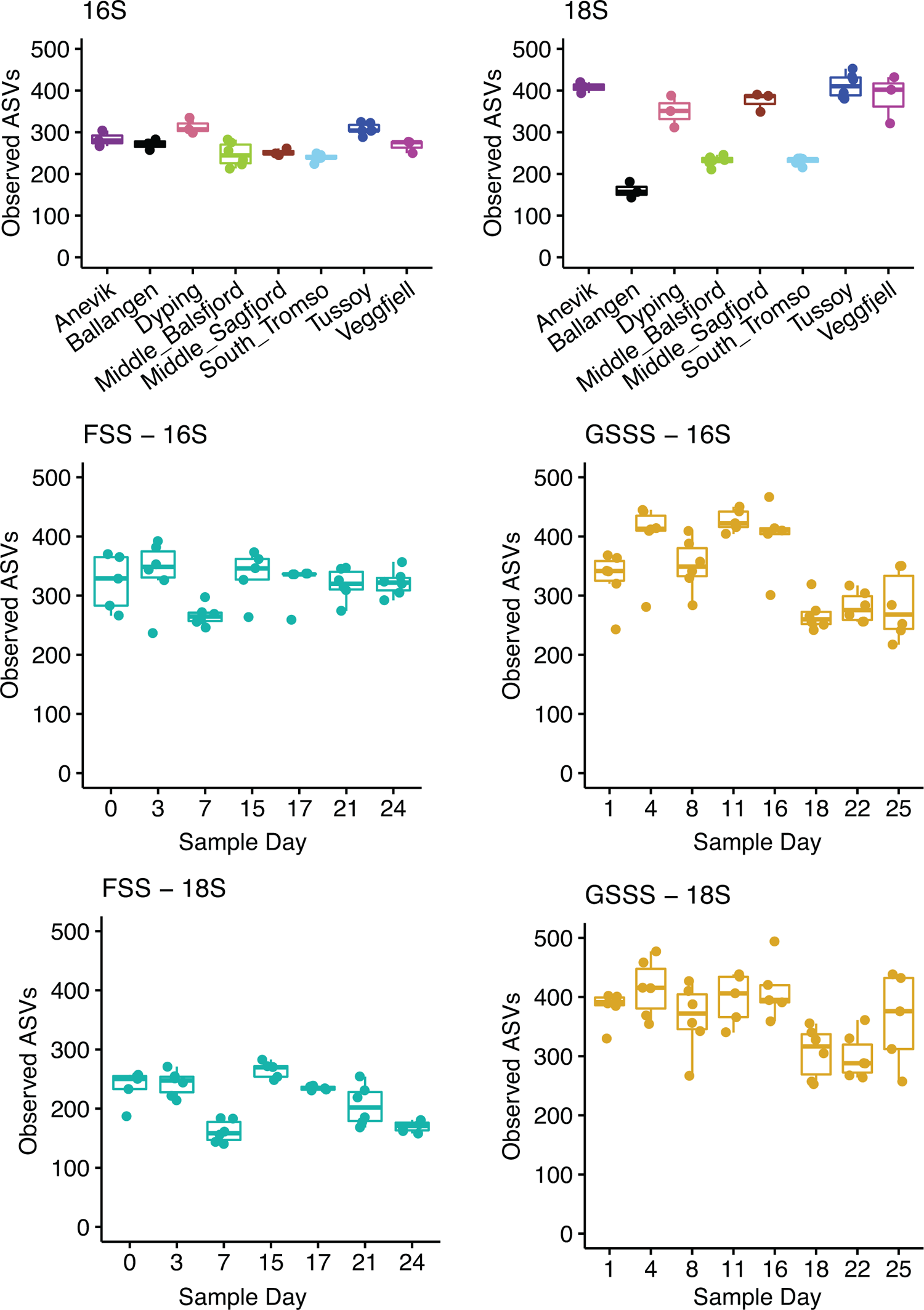
Alpha diversity as observed number of ASVs per sample set: single-time-point stations and time series stations.

---

**Supplementary Table S1.**
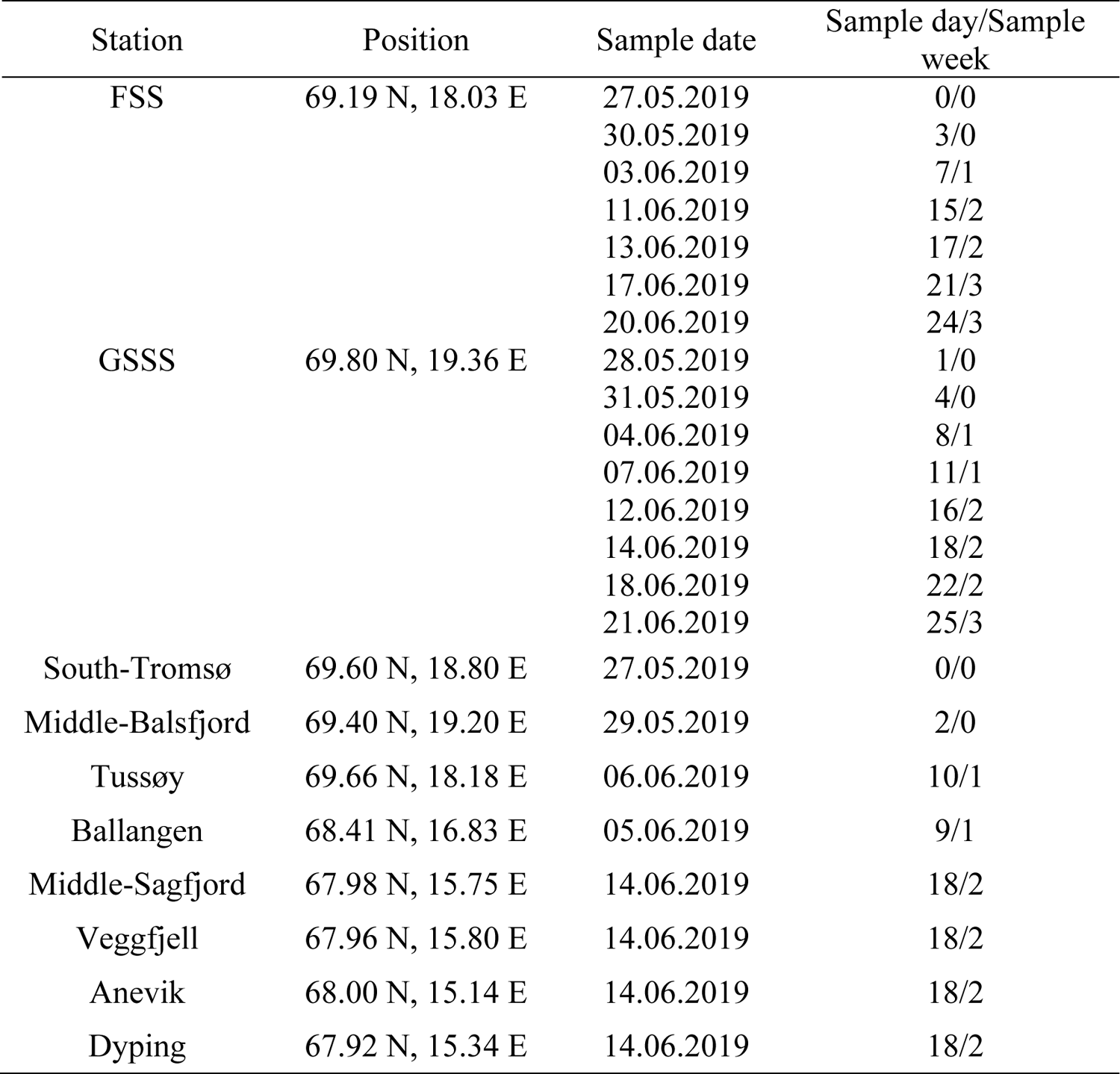
Station positions, sample dates, and corresponding sample day and week during time series sample campaign (May 27^th^ = Sample day 0 and June 21^st^ = Sample day 25).

| Station | Position | Sample date | Sample day/Sample week |
| --- | --- | --- | --- |
| FSS | 69.19 N, 18.03 E | 27.05.2019 | 0/0 |
|  |  | 30.05.2019 | 3/0 |
|  |  | 03.06.2019 | 7/1 |
|  |  | 11.06.2019 | 15/2 |
|  |  | 13.06.2019 | 17/2 |
|  |  | 17.06.2019 | 21/3 |
|  |  | 20.06.2019 | 24/3 |
| GSSS | 69.80 N, 19.36 E | 28.05.2019 | 1/0 |
|  |  | 31.05.2019 | 4/0 |
|  |  | 04.06.2019 | 8/1 |
|  |  | 07.06.2019 | 11/1 |
|  |  | 12.06.2019 | 16/2 |
|  |  | 14.06.2019 | 18/2 |
|  |  | 18.06.2019 | 22/2 |
|  |  | 21.06.2019 | 25/3 |
| South-Tromsø | 69.60 N, 18.80 E | 27.05.2019 | 0/0 |
| Middle-Balsfjord | 69.40 N, 19.20 E | 29.05.2019 | 2/0 |
| Tussøy | 69.66 N, 18.18 E | 06.06.2019 | 10/1 |
| Ballangen | 68.41 N, 16.83 E | 05.06.2019 | 9/1 |
| Middle-Sagfjord | 67.98 N, 15.75 E | 14.06.2019 | 18/2 |
| Veggfjell | 67.96 N, 15.80 E | 14.06.2019 | 18/2 |
| Anevik | 68.00 N, 15.14 E | 14.06.2019 | 18/2 |
| Dyping | 67.92 N, 15.34 E | 14.06.2019 | 18/2 |

**Supplementary Table S2.** The number of classified ASVs and sequencing depth per sample and data set before and after removal of unwanted taxa. Time series sample set includes stations FSS and GSSS; Single-time-point sample set includes stations Tussøya, South-Tromsø, Middle-Balsfjord, Ballangen, Middle-Sagfjord, Veggfjell, Anevik and Dyping; Unialgal control sample set includes *Chrysochromulina leadbeateri* UiO-035 strain culture.

| Condition | Sample set | Data set | # ASVs | Sample depth |
| --- | --- | --- | --- | --- |
| All taxa | Time series | 16S | 2761 | 54160–217118 |
|  | Time series | 18S | 7500 | 72831–34835 |
|  | Single-time-point | 16S | 1414 | 83315–168168 |
|  | Single-time-point | 18S | 3687 | 143296–469129 |
|  | Unialgal control<br>(UiO-035) | 18S | 123 | 134563–273163 |
| Removed<br>unwanted taxa | Time series | 16S | 2644 | 53962–214722 |
|  | Time series | 18S | 2674 | 56789–331887 |
|  | Single-time-point | 16S | 1368 | 83253–167400 |
|  | Single-time-point | 18S | 1682 | 118101–451910 |
|  | Unialgal control<br>(UiO-035) | 18S | 15 | 125686–265853 |

**Supplementary Table S3.** Results of PERMANOVA pariwise comparison between 1) time series sample stations, 2) sample weeks and 3) sample days within each sample station for prokaryotic and microeukaryotic communities, respectively. Unweighted UniFrac was used as a distance metric, and number of permutations was 999.

| Groups | R2 | r | p.value | Data type | Station |
| --- | --- | --- | --- | --- | --- |
| FSS vs GSSS | 0.3 | 0.54 | 0.001 | 16S |  |
| FSS vs GSSS | 0.46 | 0.68 | 0.001 | 18S |  |
| Week_0 vs Week_1 | 0.22 | 0.47 | 0.001 | 16S | FSS |
| Week_0 vs Week_2 | 0.16 | 0.39 | 0.002 | 16S | FSS |
| Week_0 vs Week_3 | 0.21 | 0.45 | 0.001 | 16S | FSS |
| Week_1 vs Week_2 | 0.25 | 0.5 | 0.001 | 16S | FSS |
| Week_1 vs Week_3 | 0.17 | 0.41 | 0.001 | 16S | FSS |
| Week_2 vs Week_3 | 0.13 | 0.36 | 0.001 | 16S | FSS |
| Week_0 vs Week_1 | 0.08 | 0.28 | 0.013 | 16S | GSSS |
| Week_0 vs Week_2 | 0.16 | 0.4 | 0.001 | 16S | GSSS |
| Week_0 vs Week_3 | 0.21 | 0.46 | 0.001 | 16S | GSSS |
| Week_1 vs Week_2 | 0.14 | 0.38 | 0.001 | 16S | GSSS |
| Week_1 vs Week_3 | 0.23 | 0.48 | 0.001 | 16S | GSSS |
| Week_2 vs Week_3 | 0.11 | 0.32 | 0.003 | 16S | GSSS |
| Week_0 vs Week_1 | 0.38 | 0.61 | 0.002 | 18S | FSS |
| Week_0 vs Week_2 | 0.33 | 0.58 | 0.001 | 18S | FSS |
| Week_0 vs Week_3 | 0.34 | 0.58 | 0.001 | 18S | FSS |
| Week_1 vs Week_2 | 0.42 | 0.65 | 0.002 | 18S | FSS |
| Week_1 vs Week_3 | 0.27 | 0.51 | 0.001 | 18S | FSS |
| Week_2 vs Week_3 | 0.24 | 0.49 | 0.001 | 18S | FSS |
| Week_0 vs Week_1 | 0.08 | 0.28 | 0.024 | 18S | GSSS |
| Week_0 vs Week_2 | 0.16 | 0.4 | 0.001 | 18S | GSSS |
| Week_0 vs Week_3 | 0.21 | 0.46 | 0.001 | 18S | GSSS |
| Week_1 vs Week_2 | 0.14 | 0.38 | 0.003 | 18S | GSSS |
| Week_1 vs Week_3 | 0.23 | 0.48 | 0.001 | 18S | GSSS |
| Week_2 vs Week_3 | 0.11 | 0.32 | 0.003 | 18S | GSSS |
| 0 vs 3 | 0.11 | 0.33 | 0.356 | 16S | FSS |
| 0 vs 7 | 0.32 | 0.57 | 0.003 | 16S | FSS |
| 0 vs 15 | 0.24 | 0.49 | 0.01 | 16S | FSS |
| 0 vs 17 | 0.27 | 0.52 | 0.006 | 16S | FSS |
| 0 vs 21 | 0.29 | 0.54 | 0.001 | 16S | FSS |
| 0 vs 24 | 0.36 | 0.6 | 0.007 | 16S | FSS |
| 3 vs 7 | 0.25 | 0.5 | 0.009 | 16S | FSS |
| 3 vs 15 | 0.21 | 0.46 | 0.006 | 16S | FSS |
| 3 vs 17 | 0.24 | 0.49 | 0.007 | 16S | FSS |
| 3 vs 21 | 0.25 | 0.5 | 0.005 | 16S | FSS |
| 3 vs 24 | 0.29 | 0.54 | 0.005 | 16S | FSS |
| 7 vs 15 | 0.36 | 0.6 | 0.004 | 16S | FSS |
| 7 vs 17 | 0.29 | 0.54 | 0.003 | 16S | FSS |
| 7 vs 21 | 0.29 | 0.54 | 0.006 | 16S | FSS |
| 7 vs 24 | 0.23 | 0.48 | 0.004 | 16S | FSS |
| 15 vs 17 | 0.22 | 0.47 | 0.009 | 16S | FSS |
| 15 vs 21 | 0.22 | 0.46 | 0.004 | 16S | FSS |
| 15 vs 24 | 0.36 | 0.6 | 0.001 | 16S | FSS |
| 17 vs 21 | 0.16 | 0.4 | 0.01 | 16S | FSS |
| 17 vs 24 | 0.27 | 0.52 | 0.003 | 16S | FSS |
| 21 vs 24 | 0.25 | 0.5 | 0.004 | 16S | FSS |
| 1 vs 4 | 0.28 | 0.53 | 0.002 | 16S | GSSS |
| 1 vs 8 | 0.23 | 0.48 | 0.004 | 16S | GSSS |
| 1 vs 11 | 0.31 | 0.56 | 0.006 | 16S | GSSS |
| 1 vs 16 | 0.34 | 0.59 | 0.004 | 16S | GSSS |
| 1 vs 18 | 0.4 | 0.63 | 0.005 | 16S | GSSS |
| 1 vs 22 | 0.29 | 0.54 | 0.002 | 16S | GSSS |
| 1 vs 25 | 0.36 | 0.6 | 0.002 | 16S | GSSS |
| 4 vs 8 | 0.18 | 0.43 | 0.002 | 16S | GSSS |
| 4 vs 11 | 0.14 | 0.38 | 0.009 | 16S | GSSS |
| 4 vs 16 | 0.17 | 0.42 | 0.002 | 16S | GSSS |
| 4 vs 18 | 0.37 | 0.61 | 0.003 | 16S | GSSS |
| 4 vs 22 | 0.31 | 0.56 | 0.004 | 16S | GSSS |
| 4 vs 25 | 0.37 | 0.61 | 0.004 | 16S | GSSS |
| 8 vs 11 | 0.21 | 0.46 | 0.001 | 16S | GSSS |
| 8 vs 16 | 0.2 | 0.44 | 0.002 | 16S | GSSS |
| 8 vs 18 | 0.33 | 0.58 | 0.002 | 16S | GSSS |
| 8 vs 22 | 0.27 | 0.52 | 0.004 | 16S | GSSS |
| 8 vs 25 | 0.32 | 0.56 | 0.004 | 16S | GSSS |
| 11 vs 16 | 0.19 | 0.43 | 0.007 | 16S | GSSS |
| 11 vs 18 | 0.41 | 0.64 | 0.002 | 16S | GSSS |
| 11 vs 22 | 0.34 | 0.58 | 0.002 | 16S | GSSS |
| 11 vs 25 | 0.38 | 0.62 | 0.002 | 16S | GSSS |
| 16 vs 18 | 0.32 | 0.56 | 0.004 | 16S | GSSS |
| 16 vs 22 | 0.27 | 0.52 | 0.001 | 16S | GSSS |
| 16 vs 25 | 0.32 | 0.56 | 0.003 | 16S | GSSS |
| 18 vs 22 | 0.15 | 0.38 | 0.005 | 16S | GSSS |
| 18 vs 25 | 0.24 | 0.49 | 0.003 | 16S | GSSS |
| 22 vs 25 | 0.18 | 0.42 | 0.004 | 16S | GSSS |
| 0 vs 3 | 0.19 | 0.43 | 0.017 | 18S | FSS |
| 0 vs 7 | 0.48 | 0.69 | 0.002 | 18S | FSS |
| 0 vs 15 | 0.5 | 0.71 | 0.008 | 18S | FSS |
| 0 vs 17 | 0.47 | 0.69 | 0.009 | 18S | FSS |
| 0 vs 21 | 0.44 | 0.67 | 0.004 | 18S | FSS |
| 0 vs 24 | 0.56 | 0.75 | 0.001 | 18S | FSS |
| 3 vs 7 | 0.42 | 0.65 | 0.002 | 18S | FSS |
| 3 vs 15 | 0.44 | 0.66 | 0.003 | 18S | FSS |
| 3 vs 17 | 0.4 | 0.63 | 0.002 | 18S | FSS |
| 3 vs 21 | 0.37 | 0.61 | 0.001 | 18S | FSS |
| 3 vs 24 | 0.49 | 0.7 | 0.004 | 18S | FSS |
| 7 vs 15 | 0.54 | 0.74 | 0.006 | 18S | FSS |
| 7 vs 17 | 0.44 | 0.66 | 0.002 | 18S | FSS |
| 7 vs 21 | 0.41 | 0.64 | 0.005 | 18S | FSS |
| 7 vs 24 | 0.37 | 0.61 | 0.002 | 18S | FSS |
| 15 vs 17 | 0.31 | 0.56 | 0.007 | 18S | FSS |
| 15 vs 21 | 0.35 | 0.59 | 0.002 | 18S | FSS |
| 15 vs 24 | 0.54 | 0.74 | 0.002 | 18S | FSS |
| 17 vs 21 | 0.23 | 0.48 | 0.001 | 18S | FSS |
| 17 vs 24 | 0.44 | 0.66 | 0.004 | 18S | FSS |
| 21 vs 24 | 0.31 | 0.55 | 0.004 | 18S | FSS |
| 1 vs 4 | 0.28 | 0.53 | 0.002 | 18S | GSSS |
| 1 vs 8 | 0.23 | 0.48 | 0.001 | 18S | GSSS |
| 1 vs 11 | 0.31 | 0.56 | 0.004 | 18S | GSSS |
| 1 vs 16 | 0.34 | 0.59 | 0.003 | 18S | GSSS |
| 1 vs 18 | 0.4 | 0.63 | 0.002 | 18S | GSSS |
| 1 vs 22 | 0.29 | 0.54 | 0.003 | 18S | GSSS |
| 1 vs 25 | 0.36 | 0.6 | 0.007 | 18S | GSSS |
| 4 vs 8 | 0.18 | 0.43 | 0.002 | 18S | GSSS |
| 4 vs 11 | 0.14 | 0.38 | 0.008 | 18S | GSSS |
| 4 vs 16 | 0.17 | 0.42 | 0.009 | 18S | GSSS |
| 4 vs 18 | 0.37 | 0.61 | 0.001 | 18S | GSSS |
| 4 vs 22 | 0.31 | 0.56 | 0.003 | 18S | GSSS |
| 4 vs 25 | 0.37 | 0.61 | 0.005 | 18S | GSSS |
| 8 vs 11 | 0.21 | 0.46 | 0.002 | 18S | GSSS |
| 8 vs 16 | 0.2 | 0.44 | 0.005 | 18S | GSSS |
| 8 vs 18 | 0.33 | 0.58 | 0.003 | 18S | GSSS |
| 8 vs 22 | 0.27 | 0.52 | 0.005 | 18S | GSSS |
| 8 vs 25 | 0.32 | 0.56 | 0.004 | 18S | GSSS |
| 11 vs 16 | 0.19 | 0.43 | 0.012 | 18S | GSSS |
| 11 vs 18 | 0.41 | 0.64 | 0.001 | 18S | GSSS |
| 11 vs 22 | 0.34 | 0.58 | 0.004 | 18S | GSSS |
| 11 vs 25 | 0.38 | 0.62 | 0.002 | 18S | GSSS |
| 16 vs 18 | 0.32 | 0.56 | 0.003 | 18S | GSSS |
| 16 vs 22 | 0.27 | 0.52 | 0.006 | 18S | GSSS |
| 16 vs 25 | 0.32 | 0.56 | 0.002 | 18S | GSSS |
| 18 vs 22 | 0.15 | 0.38 | 0.004 | 18S | GSSS |
| 18 vs 25 | 0.24 | 0.49 | 0.006 | 18S | GSSS |
| 22 vs 25 | 0.18 | 0.42 | 0.004 | 18S | GSSS |

**Supplementary Table S4.** Summary of physical environmental measurements and analyzed inorganic nutrients as well as the relative abundance of focal taxon. Mean values and (min, max) are shown.

|  | Station |  |
| --- | --- | --- |
|  | FSS | GSSS |
| Temperature (°C)<sup>a</sup> | 6.6<br>(5.4, 7.4) | 6.2<br>(5.3, 7.0) |
| Salinity<sup>a</sup> | 33.1<br>(32.5, 33.4) | 33.4<br>(33.2, 33.6) |
| MLD (m) | 3<br>(3.3) | 5<br>(3, 7) |
| Si(OH)<sub>4</sub> (μmol L<sup>-1</sup>) | 1.24<br>(0.69, 1.87) | 1.56<br>(0.09, 3.00) |
| NO<sub>2</sub><sup>-</sup> + NO<sub>3</sub><sup>-</sup> (μmol L<sup>-1</sup>) | 0.18<br>(0.07, 0.41) | 0.43<br>(0.09, 1.20) |
| PO<sub>4</sub><sup>3-</sup> (μmol L<sup>-1</sup>) | 0.06<br>(< 0.04, 0.12) | 0.07<br>(< 0.04, 0.39) |
| Focal taxon (%)<sup>b</sup> | 0.7<br>(0.4, 1.3) | 3.4<br>(0.02, 17.7) |

<sup>a</sup> Values at the depth of 10 m
<sup>b</sup> Relative abundance of the *Chrysochromulina leadbeateri* focal taxon

## REFERENCES CITED

1. Aalto, N.J., Campbell, K., Eilertsen, H.C., and Bernstein, H.C. (2021). Drivers of atmosphere-ocean CO2 flux in northern Norwegian fjords. Frontiers in Marine Science, 841. doi: 10.3389/fmars.2021.692093.

2. Aalto, N.J., Schweitzer, H., Krsmanovic, S., Campbell, K., and Bernstein, H.C. (2022). Diversity and selection of surface marine microbiomes in the Atlantic-influenced Arctic. Frontiers in Microbiology. doi: 10.3389/fmicb.2022.892634/abstract.

3. Abdi, H., and Williams, L.J. (2010). Tukey’s honestly significant difference (HSD) test. Encyclopedia of research design 3(1), 1–5.

4. Ajani, P.A., Kahlke, T., Siboni, N., Carney, R., Murray, S.A., and Seymour, J.R. (2018). The Microbiome of the Cosmopolitan Diatom Leptocylindrus Reveals Significant Spatial and Temporal Variability. Frontiers in Microbiology 9. doi: 10.3389/fmicb.2018.02758.

5. Amaral-Zettler, L.A., McCliment, E.A., Ducklow, H.W., and Huse, S.M. (2009). A method for studying protistan diversity using massively parallel sequencing of V9 hypervariable regions of small-subunit ribosomal RNA genes. PloS one 4(7), e6372.

6. Anderson, D.M., Glibert, P.M., and Burkholder, J.M. (2002). Harmful algal blooms and eutrophication: nutrient sources, composition, and consequences. Estuaries 25(4), 704–726.

7. Anderson, M.J. (2001). A new method for non-parametric multivariate analysis of variance. Austral ecology 26(1), 32–46.

8. Caporaso, J.G., Kuczynski, J., Stombaugh, J., Bittinger, K., Bushman, F.D., Costello, E.K., et al. (2010). QIIME allows analysis of high-throughput community sequencing data. Nat Methods 7(5), 335–336.

9. Coenen, A.R., Hu, S.K., Luo, E., Muratore, D., and Weitz, J.S. (2020). A primer for microbiome time-series analysis. Frontiers in genetics 11, 310.

10. Dahl, E., Bagøien, E., Edvardsen, B., and Stenseth, N.C. (2005). The dynamics of Chrysochromulina species in the Skagerrak in relation to environmental conditions. Journal of Sea Research 54(1), 15–24.

11. Davidson, K., Gowen, R.J., Tett, P., Bresnan, E., Harrison, P.J., McKinney, A., et al. (2012). Harmful algal blooms: How strong is the evidence that nutrient ratios and forms influence their occurrence? Estuarine, Coastal and Shelf Science 115, 399–413. doi: https://doi.org/10.1016/j.ecss.2012.09.019.

12. Degerlund, M., and Eilertsen, H.C. (2010). Main species characteristics of phytoplankton spring blooms in NE Atlantic and Arctic waters (68–80 N). Estuaries and coasts 33(2), 242–269.

13. Deng, Z., Chen, S., Zhang, P., Zhang, X., Adams, J.M., Luo, Q., et al. (2021). Dynamics of Free-Living and Attached Bacterial Assemblages in Skeletonema sp. Diatom Cultures at Elevated Temperatures. Frontiers in Marine Science 8. doi: 10.3389/fmars.2021.626207.

14. Du Yoo, Y., Seong, K.A., Jeong, H.J., Yih, W., Rho, J.-R., Nam, S.W., et al. (2017). Mixotrophy in the marine red-tide cryptophyte Teleaulax amphioxeia and ingestion and grazing impact of cryptophytes on natural populations of bacteria in Korean coastal waters. Harmful Algae 68, 105–117.

15. Edvardsen, B., Eikrem, W., Throndsen, J., Saez, A.G., Probert, I., and Medlin, L.K. (2011). Ribosomal DNA phylogenies and a morphological revision provide the basis for a revised taxonomy of the Prymnesiales (Haptophyta). European journal of phycology 46(3), 202–228.

16. Edvardsen, B., and Imai, I. (2006). “The ecology of harmful flagellates within Prymnesiophyceae and Raphidophyceae,” in Ecology of harmful algae. Springer), 67-79.

17. Eikrem, W., and Throndsen, J. (1998). Morphology of Chrysochromulina leadbeateri (Prymnesiophyceae) from northern Norway. Phycologia 37(4), 292–299.

18. Eilertsen, H.C., Falk-Petersen, S., Hopkins, C., and Tande, K. (1981). Ecological investigations on the plankton community of Balsfjorden, northern Norway: program for the project, study area, topography, and physical environment. Sarsia 66(1), 25–34.

19. Eilertsen, H.C., and Frantzen, S. (2007). Phytoplankton from two sub-Arctic fjords in northern Norway 2002–2004: I. Seasonal variations in chlorophyll a and bloom dynamics. Marine Biology Research 3(5), 319–332.

20. Eilertsen, H.C., and Taasen, J. (1984). Investigations on the plankton community of Balsfjorden, northern Norway. The phytoplankton 1976–1978. Environmental factors, dynamics of growth, and primary production. Sarsia 69(1), 1-15.

21. Estep, K. (1984). Chloroplast containing microflagellates in natural populations of north Atlantic nanoplankton, their identification and distribution; including a description of 5 new species of Chrysochromulina (Prymnesiophyceae). Protistologica 20, 613–634.

22. Fuhrman, J.A., Cram, J.A., and Needham, D.M. (2015). Marine microbial community dynamics and their ecological interpretation. Nature Reviews Microbiology 13(3), 133–146.

23. Gilbert, J.A., Meyer, F., Antonopoulos, D., Balaji, P., Brown, C.T., Brown, C.T., et al. (2010). Meeting report: the terabase metagenomics workshop and the vision of an Earth microbiome project. Stand Genomic Sci 3(3), 243.

24. Grann-Meyer, E. (2020). Chrysochromulina leadbeateri - Understanding the presumed causal agent behind the harmful algal bloom of 2019. Master’s thesis, UiT - Arctic University of Norway.

25. Guillard, R.R. (1975). “Culture of phytoplankton for feeding marine invertebrates,” in Culture of marine invertebrate animals. Springer), 29-60.

26. Hallegraeff, G.M., Anderson, D.M., Belin, C., Bottein, M.-Y.D., Bresnan, E., Chinain, M., et al. (2021). Perceived global increase in algal blooms is attributable to intensified monitoring and emerging bloom impacts. Communications Earth & Environment 2(1), 1–10.

27. Hegseth, E.N., and Eilertsen, H.C. (1991). The Chrysochromulina leadbeateri bloom in Troms May/June 1991. Development and causes.. Fisk. Hav. 3, 45–61.

28. Heidal, K., and Mohus, A. (1995). The toxic Chrysochromulina--salmon disaster of 1991 in northern Norway with some follow-up monitoring records of 1992 and-93. *LAVOISIER*, PARIS(FRANCE). 163–168.

29. John, U., Šupraha, L., Gran-Stadniczeñko, S., Bunse, C., Cembella, A., Eikrem, W., et al. (2022). Spatial and biological oceanographic insights into the massive fish-killing bloom of the haptophyte Chrysochromulina leadbeateri in northern Norway. Harmful Algae, 102287.

30. Karlsen, K.M., Robertsen, R., and Hersoug, B. (2019). Mapping the course of events and preparedness for algal blooms in spring 2019 - Astafjorden, Ofotfjorden, Vestfjorden og Tysfjorden. Nofima AS 29, 1–44.

31. Karlson, B., Andersen, P., Arneborg, L., Cembella, A., Eikrem, W., John, U., et al. (2021). Harmful algal blooms and their effects in coastal seas of Northern Europe. Harmful Algae 102, 101989.

32. Kaufman, L., and Rousseeuw, P.J. (2009). Finding groups in data: an introduction to cluster analysis. New York: John Wiley & Sons.

33. Lindh, M.V., Sjöstedt, J., Andersson, A.F., Baltar, F., Hugerth, L.W., Lundin, D., et al. (2015). Disentangling seasonal bacterioplankton population dynamics by high-frequency sampling. Environmental microbiology 17(7), 2459–2476.

34. Liu, C., Cui, Y., Li, X., and Yao, M. (2021). microeco: an R package for data mining in microbial community ecology. FEMS Microbiology Ecology 97(2), fiaa255.

35. Lord, E., Willems, M., Lapointe, F.-J., and Makarenkov, V. (2017). Using the stability of objects to determine the number of clusters in datasets. Information Sciences 393, 29–46.

36. Love, M.I., Huber, W., and Anders, S. (2014). Moderated estimation of fold change and dispersion for RNA-seq data with DESeq2. Genome biology 15(12), 1–21.

37. Lozupone, C., Lladser, M.E., Knights, D., Stombaugh, J., and Knight, R. (2011). UniFrac: an effective distance metric for microbial community comparison. The ISME journal 5(2), 169–172.

38. Martin-Platero, A.M., Cleary, B., Kauffman, K., Preheim, S.P., McGillicuddy, D.J., Alm, E.J., et al. (2018). High resolution time series reveals cohesive but short-lived communities in coastal plankton. Nature communications 9(1), 1–11.

39. Massana, R., Terrado, R., Forn, I., Lovejoy, C., and Pedrós-Alió, C. (2006). Distribution and abundance of uncultured heterotrophic flagellates in the world oceans. Environmental microbiology 8(9), 1515–1522.

40. Needham, D.M., and Fuhrman, J.A. (2016). Pronounced daily succession of phytoplankton, archaea and bacteria following a spring bloom. Nature Microbiology 1(4), 16005. doi: 10.1038/nmicrobiol.2016.5.

41. Nielsen, T.G., Kiørboe, T., and Bjørnsen, P.K. (1990). Effects of a Chrysochromulina polylepis subsurface bloom on the planktonic community. Marine Ecology Progress Series, 21-35.

42. Oksanen, J., Blanchet, F.G., Kindt, R., Legendre, P., Minchin, P.R., O’hara, R., et al. (2013). Package ‘vegan’. Community ecology package, version 2(9).

43. Parte, A.C., Carbasse, J.S., Meier-Kolthoff, J.P., Reimer, L.C., and Göker, M. (2020). List of Prokaryotic names with Standing in Nomenclature (LPSN) moves to the DSMZ. International Journal of Systematic and Evolutionary Microbiology 70(11), 5607.

44. Peralta-Ferriz, C., and Woodgate, R.A. (2015). Seasonal and interannual variability of pan-Arctic surface mixed layer properties from 1979 to 2012 from hydrographic data, and the dominance of stratification for multiyear mixed layer depth shoaling. Progress in Oceanography 134, 19–53.

45. R Core Team (2021). R: A language and environment for statistical computing, R Foundation for Statistical Computing.

46. Reigstad, M., and Wassmann, P. (1996). Importance of advection for pelagic-benthic coupling in north Norwegian fjords. Sarsia 80(4), 245–257.

47. Rey, F. (1991). The Chrysochromulina leadbeateri bloom in Vestfjorden, north Norway, May-June 1991: Proceedings from a scientific working meeting. Fisk. Hav. 3, 1-122.

48. Rey, F., and Aure, J. (1991). The Chrysochromulina leadbeateri bloom in the Vestfjord, north Norway, May-June 1991. Environmental conditions and possible causes. Fisk. Hav. 3, 13-32.

49. Statistics Norway (2020). Aquaculture 2019 [Online]. Statistics Norway. Available: https://www.ssb.no/jord-skog-jakt-og-fiskeri/statistikker/fiskeoppdrett [Accessed 23.04 2022].

50. Teeling, H., Fuchs, B.M., Bennke, C.M., Krueger, K., Chafee, M., Kappelmann, L., et al. (2016). Recurring patterns in bacterioplankton dynamics during coastal spring algae blooms. *elife* 5, e11888.

51. Uronen, P., Lehtinen, S., Legrand, C., Kuuppo, P., and Tamminen, T. (2005). Haemolytic activity and allelopathy of the haptophyte Prymnesium parvum in nutrient-limited and balanced growth conditions. Marine Ecology Progress Series 299, 137–148.

52. Wassmann, P., Svendsen, H., Keck, A., and Reigstad, M. (1996). Selected aspects of the physical oceanography and particle fluxes in fjords of northern Norway. Journal of Marine Systems 8(1-2), 53–71.

53. Zhou, J., Richlen, M.L., Sehein, T.R., Kulis, D.M., Anderson, D.M., and Cai, Z. (2018). Microbial Community Structure and Associations During a Marine Dinoflagellate Bloom. Frontiers in Microbiology 9. doi: 10.3389/fmicb.2018.01201.

